# Functional characterisation of banana (*Musa* spp.) 2-oxoglutarate-dependent dioxygenases involved in flavonoid biosynthesis

**DOI:** 10.1101/2021.05.03.442406

**Authors:** Mareike Busche, Christopher Acatay, Stefan Martens, Bernd Weisshaar, Ralf Stracke

## Abstract

Bananas (*Musa*) are non-grass, monocotyledonous, perennial plants that are well-known for their edible fruits. Their cultivation provides food security and employment opportunities in many countries. Banana fruits contain high levels of minerals and phytochemicals, including flavonoids, which are beneficial for human nutrition. To broaden the knowledge on flavonoid biosynthesis in this major crop plant, we aimed to identify and functionally characterise selected structural genes encoding 2-oxoglutarate-dependent dioxygenases, involved in the formation of the flavonoid aglycon.

*Musa* candidates genes predicted to encode flavanone 3-hydroxylase (F3H), flavonol synthase (FLS) and anthocyanidin synthase (ANS) were assayed. Enzymatic functionalities of the recombinant proteins were confirmed *in vivo* using bioconversion assays. Moreover, transgenic analyses in corresponding *Arabidopsis thaliana* mutants showed that *MusaF3H, MusaFLS* and *MusaANS* were able to complement the respective loss-of-function phenotypes, thus verifying functionality of the enzymes *in planta*.

Knowledge gained from this work provides a new aspect for further research towards genetic engineering of flavonoid biosynthesis in banana fruits to increase their antioxidant activity and nutritional value.

## Introduction

Banana (*Musa* spp.) plants are well-known for their edible fruit and serve as a staple food crop in Africa, Central and South America (Arias et al., 2003). With more than 112 million tons produced in 2016, bananas are among the most popular fruits in the world and provide many employment opportunities (FAO, 2019). Furthermore, banana fruits are rich in health promoting minerals and phytochemicals, including flavonoids, a class of plant specialised metabolites, which contribute to the beneficial effects through their antioxidant characteristics (Forster et al., 2003; Wall, 2006; Singh et al., 2016). Flavonoid molecules share a C6-C3-C6 aglycon core, which can be reorganised or modified, e.g., by oxidation or glycosylation (Tanaka et al., 2008; Le Roy et al., 2016). Modifications at the central ring structure divide flavonoids into ten major subgroups (i.e. chalcones, aurones, flavanones, flavones, isoflavones, dihydroflavonols, flavonols, leucoanthocyanidins, anthocyanidins, and flavan-3-ols). The diversity in chemical structure is closely related to diverse bioactivities of flavonoids in plant biology and human nutrition (Falcone Ferreyra et al., 2012). For example, anthocyanins are in many cases well known to colour flowers and fruits to attract animals and thus promoting pollination and dispersion of seeds (Ishikura and Yoshitama, 1984; Gronquist et al., 2001; Grotewold, 2006). Flavonols can interact with anthocyanins to modify the colour of fruits (Andersen and Jordheim, 2010) and play a prominent role in protection against UV-B irradiation (Li et al., 1993) and also in plant fertility (Mo et al., 1992).

Many researchers have attributed positive effects on human health to flavonoids: e.g., antigenotoxic (Dauer et al., 2003), anticarcinogenic and antioxidative (Kandil et al., 2002) effects, as well as the prevention of cardiovascular diseases have been suggested (summarised in Perez-Vizcaino and Duarte, 2010). Additionally, Sun et al. (2019) suggested an involvement of flavonoids in the plant defence against the tropical race 4 (TR4) of the *Musa* Fusarium wilt (commonly known as ‘panama disease’) pathogen *Fusarium oxysporum* f. sp. *cubense* (Foc), which is a threat to the global banana production. It is certainly interesting to take a closer look at the biosynthesis of flavonoids in *Musa*.

Flavonoids are derived from the amino acid L-phenylalanine and malonyl-coenzyme A. Their biosynthesis (Figure 1) has been analysed in many different species including the dicotyledonous model plant *Arabidopsis thaliana* (*A. thaliana*) (Hahlbrock and Scheel, 1989; Lepiniec et al., 2006) and the monocotyledonous crop plants *Zea mays* (*Z. mays*) (summarised in Tohge et al., 2017) and *Oryza sativa* (*O. sativa*) (summarised in Goufo and Trindade, 2014). The first committed step, catalysed by the enzyme chalcone synthase (CHS), is the formation of naringenin chalcone from *p*-coumaroyl CoA and malonyl CoA (Kreuzaler and Hahlbrock, 1972). A heterocyclic ring is introduced during the formation of naringenin (a flavanone) from naringenin chalcone, which can occur spontaneously or catalysed by chalcone isomerase (CHI) (Bednar and Hadcock, 1988). The enzyme flavanone 3-hydroxylase (F3H or FHT) converts flavanones to dihydroflavonols by hydroxylation at the C-3 position (Forkmann et al., 1980). Alternatively, flavanones can be converted to flavones by flavone synthase I or II (FNSI or II) (Britsch, 1990). Flavonol synthase (FLS) catalyses the conversion of dihydroflavonols to the corresponding flavonols by introducing a double bond between C-2 and C-3 (Forkmann et al., 1986; Holton et al., 1993). Moreover, dihydroflavonols can be converted to leucoanthocyanidins by dihydroflavonol 4-reductase (DFR) (Heller et al., 1985), which competes with FLS for substrates (Luo et al., 2016). Anthocyanidin synthase (ANS, also termed leucoanthocyanidin dioxygenase, LDOX) converts leucoanthocyanidins to anthocyanidins (Saito et al., 1999).

**Figure 1:**
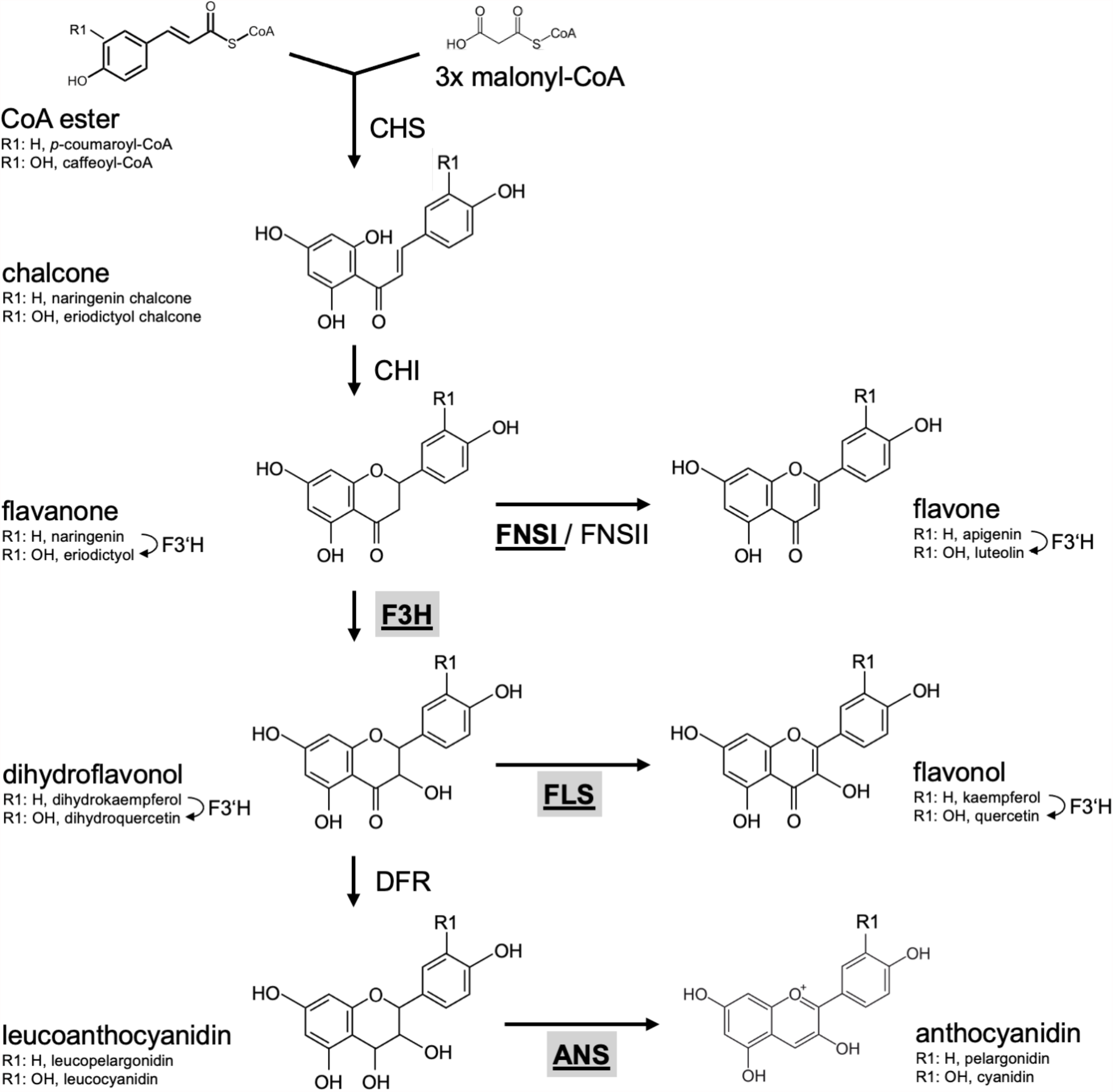
Schematic, simplified illustration of the core flavonoid aglycon biosynthesis pathway in plants. Chalcone synthase (CHS), the first committed enzyme in flavonoid biosynthesis, connects a CoA ester and malonyl-CoA, forming a chalcone. Chalcone isomerase (CHI) introduces the heterocyclic ring. The resulting flavanone is converted to dihydroflavonol by flavanone 3-hydroxylase (F3H) or to flavones by flavone synthase (FNSI or FNSII). Dihydroflavonols are converted to flavonols by flavonol synthase (FLS). Alternatively, dihydroflavonol 4-reductase (DFR) can reduce dihydroflavonols to the corresponding leucoanthocyanidin, which is converted to anthocyanidin by the activity of anthocyanidin synthase (ANS). Flavonoid 3’ hydroxylase (F3’H) hydroxylates 3’ position of the B-ring using different flavonoid substrates. 2-oxoglutarate-dependant oxygenases are given in bold underlined. Enzymes in the focus of this work are highlighted by grey boxes.

The flavonoid biosynthesis enzymes belong to different functional classes (summarised in Winkel, 2006): polyketide synthases (e.g., CHS), 2-oxoglutarate-dependent dioxygenases (2-ODD, e.g., F3H, ANS, FLS, FNSI), short-chain dehydrogenases/reductases (e.g., DFR), aldo-keto reductases (e.g., chalcone reductase, CHR) and cytochrome P450 monooxygenases (e.g., FNSII, F3′H). Flavonoid biosynthesis is evolutionary old in plants (Wen et al., 2020) and, although differences exist, quite similar in dicots like *A. thaliana* and monocots like *Musa*.

In the present study, we focus on the functional class of flavonoid aglycon forming enzymes, namely the 2-ODDs. 2-ODDs occur throughout the kingdom of life and play important roles in many biological processes including oxygen sensing and DNA repair (Jaakkola et al., 2001; Trewick et al., 2002). They are a class of non-heme iron-containing enzymes, which require 2-oxoglutarate, Fe^2+^ and ascorbate for substrate conversion (summarised in Prescott and John, 1996).

Plant 2-ODDs share conserved amino acid residues which coordinate ferrous iron binding (HxDxnH) and binding of 2-oxoglutarate (RxS) (Cheng et al., 2014). They can be divided into three distinct evolutionary classes, DOXA, DOXB and DOXC, based on amino acid similarity (Kawai et al., 2014). The 2-ODDs involved in specialised metabolism were classified into the DOXC class. Yet, all flavonoid biosynthesis related 2-ODDs were found in the DOXC subclades 28 (F3H and FNSI) or 47 (FLS and ANS) (Kawai et al., 2014). In some cases, the high amino acid similarity between the different enzymes leads to overlapping functions. For example, FLSs from *Ginkgo biloba* (*G. biloba*) (Xu et al., 2012) and *O. sativa* (Park et al., 2019) can accept flavanones and dihydroflavonols as substrates. Also, ANSs from *A. thaliana* (Stracke et al., 2009) and *Malus domestica* (*M. domestica*) (Yan et al., 2005) can catalyse the formation of flavonol glycosides.

Based on the *Musa acuminata* (*M. acuminata*) genome sequence annotation (DHont et al., 2012), Pandey et al. (2016) identified 28 putative *Musa* flavonoid biosynthesis enzymes. This includes seven 2-ODD-type enzymes, namely two F3H, four FLS and one ANS, while no suitable FNSI candidate was identified. While the expression of the respective genes and correlations of expression to flavonoid metabolite accumulation have been studied, the functionality of these enzymes has not been analysed until now.

Here, we describe the sequence-based and functional characterisation of 2-ODDs from the non-grass monocot *Musa*. Functionalities of recombinant *Musa* enzymes were analysed using *in vivo* bioconversion assays and by *in planta* complementation of corresponding *A. thaliana* loss-of-function mutants. This resulted in the experimental confirmation that the *Musa* flavonoid 2-ODDs studied have the predicted functions. The presented results contribute to the understanding of flavonoid biosynthesis in *Musa*. They provide a strong basis for further research to enhance the efficiency of flavonoid production in banana, in order to increase the fruits’ health promoting effects.

## Materials and Methods

### Plant material

Banana plants (Grand Naine, plantain) for RNA extraction were grown in the field in Lucknow, India. *Musa* gene annotation identifiers refer to the study of Martin et al. (2016). Columbia-0 (Col-0, NASC ID N1092) and Nössen-0 (Nö-0, NASC ID N3081) were used as wildtype controls. The *A. thaliana* mutants *tt6-2* (*f3h*, GK-292E08, Col-0 background) (Appelhagen et al., 2014) and *ans/fls1-2* (synonym *ldox/fls1-2, ldox: SALK_028793*, Col-0 background; *fls1-2:* RIKEN_PST16145, Nö-0 background) (Stracke et al., 2009) were used for complementation experiments.

### Phylogenetic analysis

Multiple protein sequence alignments were created with MAFFT v7 (Katoh and Standley, 2013) using default settings. The Approximately-Maximum-Likelihood phylogenetic tree of 33 plant 2-ODDs with proven F3H, FNSI, FLS and ANS functionality (Supplementary Table S1) and seven *Musa* 2-ODDs was constructed as described by Pucker et al. (2020b): MAFFT alignments were cleaned with pxclsq (Brown et al., 2017) and the tree was constructed with FastTree v2.1.10 using the WAG+CAT model (Price et al., 2010). The tree was visualised with Interactive Tree Of Life (Letunic and Bork, 2019), branch lengths were neglected.

### Expression analysis

Expression data for *Musa* 2-ODD genes was extracted from previous analyses (Pucker et al., 2020a). Short Read Archive IDs can be found in Supplementary Table S2.

### Total RNA extraction, cDNA synthesis and molecular cloning

Isolation of RNA from different *Musa* plant organs (leaf, pseudostem, bract, fruit peel, fruit pulp) was performed according to a protocol from Asif et al. (2000). cDNA synthesis was performed from 1 µg total RNA using the ProtoScript^®^ First Strand cDNA Synthesis Kit (New England Biolabs, NEB) with the provided random primer mix according to the suppliers’ instructions. Amplification of predicted full length CDSs (Pandey et al., 2016) were done using Q5® High-Fidelity DNA polymerase (NEB) and gene-specific primers (Supplementary Table S3) according to standard protocols. Creation of full-length coding sequence constructs (CDS) was performed using the GATEWAY^®^ Technology (Invitrogen). *MusaF3H1* (Ma02_t04650), *MusaF3H2* (Ma07_t17200), *MusaFLS1* (Ma03_t06970), *MusaFLS3* (Ma10_t25110) and *MusaANS* (Ma05_t03850) CDSs were successfully amplified on different cDNA pools. The resulting PCR products were recombined into pDONR™/Zeo (Invitrogen) with BP clonase (Invitrogen) resulting in Entry plasmids, which were sequenced by Sanger technology (Sanger et al., 1977) on 3730XL sequencers using BigDye terminator v3.1 chemistry (Thermo Fisher). Entry plasmids for *AtF3H, AtFLS1* and *AtMYB12* were available from previous studies (Preuss et al., 2009; Stracke et al., 2017), *PcFNSI* was amplified on a plasmid from a previous study (Martens et al., 2003). The full-length CDSs were introduced from the Entry plasmids into the inducible *E. coli* expression vector pDEST17 (Invitrogen) and the binary expression vector pLEELA (Jakoby et al., 2004) using GATEWAY LR-reaction (Invitrogen).

### Heterologous expression in *E. coli*

pDEST17-based plasmids containing *proT7*-RBS-6xHis-CDS-*T7term* expression cassettes (*proT7: T7 promoter*, RBS: ribosome binding site, 6xHis: polyhistidine tag, *T7term: T7 transcription terminator*) were transformed into BL21-AI cells (Invitrogen). Cultures were grown in LB to an OD_600_ of about 0.4 and expression was induced with 0.2 % L-arabinose.

### F3H and FLS bioconversion assay in *E. coli*

The enzyme assay was performed using 20 mL *E. coli* cultures expressing the respective constructs right after induction with L-arabinose. 100 µL substrate (10 mg/mL naringenin, eriodictyol or dihydroquercetin), 50 µL 2-oxoglutaric acid, 50 µL FeSo_4_ and 50 µL 1 M sodium ascorbate were added. The cultures were incubated at 28 °C overnight. To extract flavonoids, 1 mL was removed from each culture and mixed with 200 µL ethyl acetate by vortexing for 30 s. After centrifugation for 2 min. at 14,000 g, the organic phase was transferred into a fresh reaction tube. Samples were taken after 0 h, 1 h, 2 h, 3 h, 4 h and 24 h. Flavonoid content was analysed by high-performance thin-layer chromatography (HPTLC). Naringenin (Sigma), dihydrokaempferol (Sigma), kaempferol (Roth), eriodictyol (TransMIT PlantMetaChem), apigenin (TransMIT PlantMetaChem), dihydroquercetin (Roth) and quercetin (Sigma) were dissolved in methanol and used as standards. 3 µL of each methanolic extract were spotted on a HPTLC Silica Gel 60 plate (Merck). The mobile phase was composed of 50 % chloroform, 45 % acetic acid and 5 % water. Flavonoid compounds were detected as described previously (Stracke et al., 2007), using diphenylboric acid 2-aminoethyl ester (DPBA) and UV-light (Sheahan and Rechnitz, 1992).

### *Agrobacterium*-mediated transformation of *A. thaliana*

T-DNA from pLEELA-based plasmids containing *2xpro35S*-driven *Musa2-ODDs*, were transformed into *A. thaliana* plants via *Agrobacterium tumefaciens* (Agrobacterium, GV101::pMP90RK, (Koncz and Schell, 1986)) mediated gene transfer using the floral dip method (Clough and Bent, 1998). Successful T-DNA integration was verified by BASTA selection and PCR-based genotyping.

### Flavonoid staining of *A. thaliana* seedlings

*In situ* visualisation of flavonoids in norflurazon-bleached *A. thaliana* seedlings was performed according to Stracke et al. (2007) using DPBA/Triton X-100 and epifuorescence microscopy.

### Analysis of flavonols in methanolic extracts

Flavonol glycosides were extracted and analysed as previously described (Stracke et al., 2009). *A. thaliana* rosette leaves were homogenised in 80 % methanol, incubated for 15 min. at 70 °C and centrifuged for 10 min. at 14,000 g. Supernatants were vacuum-dried. The dried pellets were dissolved in 80 % methanol and analysed by HPTLC on a Silica Gel 60 plate (Merck). Ethyl acetate, formic acid, acetic acid, water (100:26:12:12) were used as a mobile phase. Flavonoid compounds were detected as described above.

### Determination of anthocyanin content

To induce anthocyanin production, *A. thaliana* seedlings were grown on 0.5 MS plates with 4 % sucrose and 16 h of light illumination per day at 22 °C. Six day old seedlings were used to photometrically quantify anthocyanins as described by Mehrtens et al. (2005). All samples were measured in three independent biological replicates. Error bars indicate the standard error of the average anthocyanin content. Statistical analysis was performed using the Mann– Whitney U test (Mann and Whitney, 1947).

## Results

### Creation of cDNA constructs and sequence based characterisation of *Musa* flavonoid 2-ODDs

Previously described putative *Musa* flavonoid biosynthesis enzymes encoded in the *M. acuminata* Pahang DH reference genome sequence were used. This included two F3Hs, four FLSs and one ANS, which were classified as 2-ODDs. To functionally characterise these *Musa* enzymes, we amplified the corresponding CDSs from a cDNA template collection derived from different *Musa* organs, using primers designed on the Pahang DH reference genome sequence. The successfully amplified cDNAs of *MusaF3H1* and *MusaF3H2* were derived from plantain pulp, *MusaFLS1* from Grand Naine young leaf, *MusaFLS3* on Grand Naine bract, and *MusaANS* on Grand Naine peel. Unfortunately, we were not able to amplify *MusaFLS2* and *MusaFLS4* from our template collection.

Comparison of the resulting 2-ODD cDNA sequences with the reference sequence revealed several single-nucleotide polymorphisms (SNPs) (Supplementary File S1). The derived amino acid sequences show close similarity to other plant 2-ODD proteins known to be involved in flavonoid biosynthesis (Supplementary File S2). The amino acids well known to coordinate ferrous iron binding (HxDxnH) and binding of 2-oxoglutarate (RxS) are also conserved in the *Musa* 2-ODDs. Additionally, the *At*FLS1 residues which have been shown to be involved in flavonoid substrate binding (Supplementary File S2) are conserved. The residue F293 (all positions refer to *At*FLS1) is conserved in all *Musa* 2-ODD proteins, F134 and K202 are found in the *Musa*FLSs and *Musa*ANS and E295 is conserved in *Musa*ANS.

To analyse the evolutionary relationship between *Musa* 2-ODDs and 33 known flavonoid biosynthesis related 2-ODDs, a phylogenetic tree was built (Figure 2). The phylogenetic tree revealed two distinct clades, which correspond to the DOXC28 and DOXC47 classes of the 2-ODD superfamily. In the F3H and FNSI-containing DOXC28 class, *Musa*F3H1 and *Musa*F3H2 cluster with other F3Hs from monocotyledonous plants. In the FLS and ANS-containing DOXC47 class, the *Musa*ANS clusters with ANSs from monocotyledonous plants, while the FLSs from monocotyledonous plants do not form a distinct group, although the *Mus*aFLSs are in proximity to *Zm*FLS and *Os*FLS.

**Figure 2:**
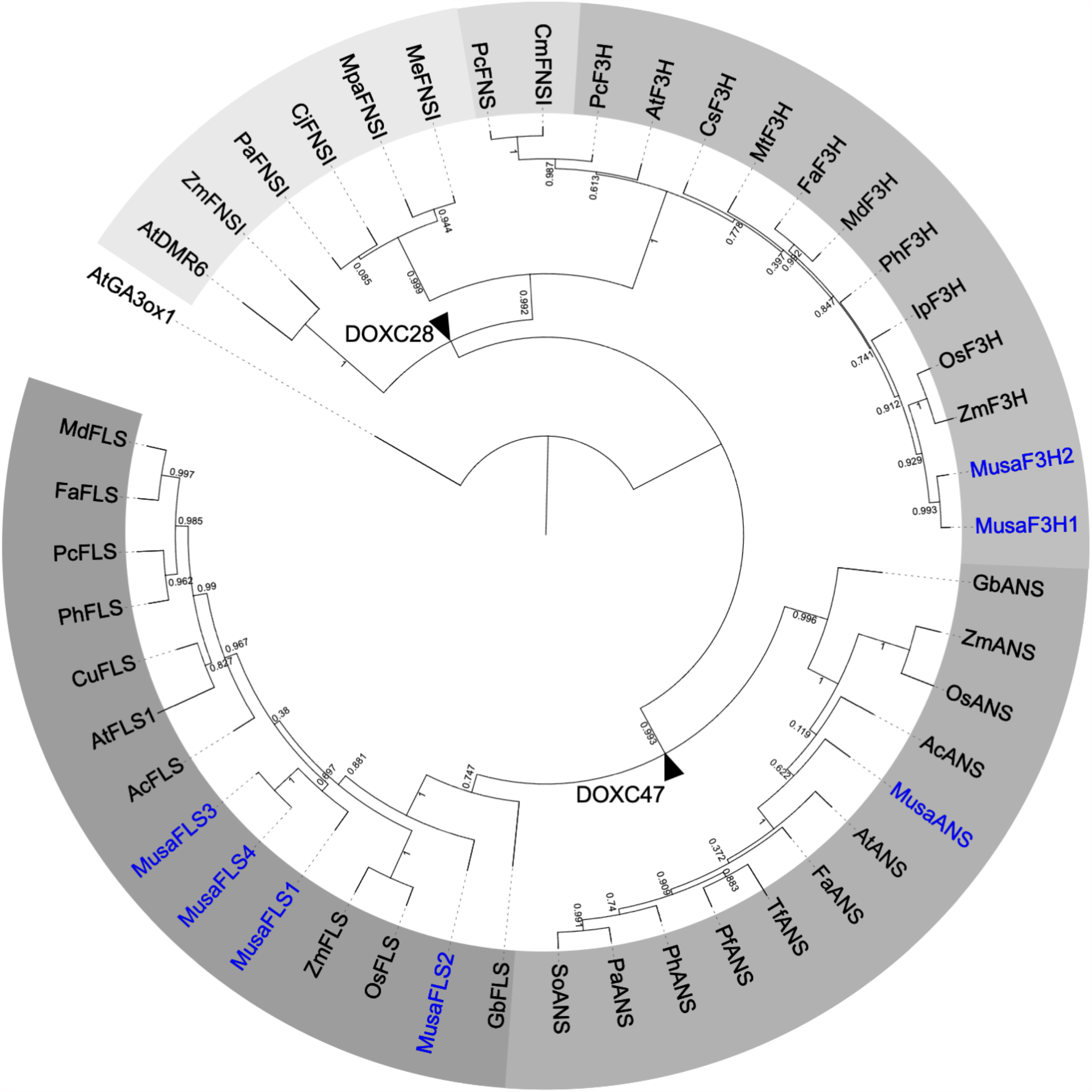
Rooted Approximately-Maximum-Likelihood (ML) phylogenetic tree of 2-ODDs involved in flavonoid biosynthesis. ODDs from Banana are given in blue, 39 enzymes with proven F3H, FNSI, FLS or ANS activity were included. Different grey scales indicate F3H, FLS, ANS and two evolutionary FNSI clades. *At*GA3ox1 (gibberellin 3 beta-hydroxylase1) was used as an outgroup. Branchpoints to DOXC28 and DOXC47 classes are marked with black arrowheads.

Our analyses clearly show that the CDSs identified for *Musa*F3H1, *Musa*F3H2, *Musa*FLS1, *Musa*FLS3 and *Musa*ANS have the potential to encode functional flavonoid 2-ODDs.

### Expression profiles of *Musa 2-ODD* genes

Analysis of the expression patterns of the genes studied was performed using published RNA-Seq data. We obtained normalised RNA-Seq read values for *Musa* 2-ODD genes from several organs and developmental stages (Table 1). *MusaF3H1* and *MusaF3H2* are expressed in almost all analysed organs and developmental stages. *MusaF3H1* expression is highest in early developmental stages of pulp (S1, S2), followed by intermediate developmental stages of peel (S2, S3). *MusaF3H2* shows highest expression in adult leaves. *MusaFLS2* was not expressed in any of the analysed samples. All four *MusaFLS* genes show low or no expression in seedlings and embryonic cell suspension. *MusaFLS1* shows variance in transcript abundance with particularly high levels in peel (S2) and in young and adult leaves. *MusaFLS3* and *MusaFLS4* transcript levels are comparatively constant and low, with highest transcript abundance in root and peel (S1 and S3 respectively). While *MusaFLS1* transcript abundance is highest in adult leaf, *MusaFLS3* and *MusaFLS4* lack expression in this tissue. *MusaANS* shows highest transcript abundance in pulp (S1, S2, S4) and peel (S3) and very low expression in embryogenic cell suspension, seedlings and leaves.

**Table 1:** Expression profiles of *Musa2-*ODD genes in different organs and developmental stages based on RNA-Seq data. Expression levels are given in fragments per kilo base per million mapped reads (FPKM). Different grey scales indicate expression levels with higher levels given in darker grey. S1-S4 indicate stages of pulp and peel developments from immature to ripe (Lu et al., 2018).

| gene | embryogenic | seedling | root | leaf |  |  |  | pulp |  |  |  |  | peel |  |  |  |  |
| --- | --- | --- | --- | --- | --- | --- | --- | --- | --- | --- | --- | --- | --- | --- | --- | --- | --- |
|  |  |  |  | leaf | young leaf | adult leaf | old leaf | pulp | pulp S1 | pulp S2 | pulp S3 | pulp S4 | peel | peel S1 | peel S2 | peel S3 | peel S4 |
| *MusaF3H1* | 3 | 9 | 80 | 44 | 73 | 27 | 3 | 170 | 385 | 191 | 43 | 60 | 82 | 46 | 140 | 104 | 36 |
| *MusaF3H2* | 21 | 0 | 23 | 57 | 39 | 171 | 7 | 32 | 96 | 14 | 3 | 14 | 19 | 7 | 27 | 33 | 9 |
| *MusaFLS1* | 6 | 0 | 4 | 189 | 91 | 643 | 16 | 2 | 2 | 1 | 2 | 2 | 29 | 3 | 99 | 7 | 7 |
| *MusaFLS2* | 0 | 0 | 0 | 0 | 0 | 0 | 0 | 0 | 0 | 0 | 0 | 0 | 0 | 0 | 0 | 0 | 0 |
| *MusaFLS3* | 0 | 0 | 32 | 16 | 17 | 0 | 0 | 16 | 15 | 22 | 10 | 17 | 22 | 27 | 16 | 20 | 25 |
| *MusaFLS4* | 0 | 0 | 19 | 4 | 0 | 0 | 0 | 10 | 11 | 4 | 12 | 13 | 15 | 16 | 11 | 19 | 14 |
| *MusaANS* | 1 | 3 | 90 | 41 | 79 | 3 | 1 | 255 | 559 | 298 | 26 | 137 | 73 | 47 | 50 | 163 | 30 |

### *Musa*F3H1 and *Musa*F3H2 are functional flavanone 3-hydroxylases

To confirm F3H activity *in vivo*, we used a bioconversion assay. *MusaF3H1* and *MusaF3H2* were heterologously expressed in *E. coli* and the bacterial cultures were fed with naringenin or eriodictyol as substrate of F3H. Both recombinant proteins, *Musa*F3H1 and *Musa*F3H2, were able to convert naringenin to dihydrokaempferol (DHK) in the presence of 2-oxoglutarate and ferrous iron (Figure 3A). After 24 h of incubation, only small amounts of naringenin remained un-converted. Conversion of the formed DHK to kaempferol by *Musa*F3H1 or *Musa*F3H2 was not observed. Furthermore, eriodictyol was converted to dihydroquercetin (DHQ) by *Musa*F3H1 and *Musa*F3H2 (Supplementary File S3). A further conversion to the quercetin was not observed. Since *Musa*F3H2 was previously considered as FNSI candidate, we analysed FNSI activity in a bioconversion assay. While *Musa*F3H2 was able to convert naringenin to DHK we were not able to detect apigenin product in our approach (Supplementary File S4). Accordingly, *Musa*F3H2 does not show FNSI activity in our assay. For further *in planta* analysis we chose a complementation assay with an *A. thaliana f3h* mutant *(tt6-2)* which expresses *FLS* and *ANS* but no *F3H. MusaF3H1* or *MusaF3H2* were expressed in *tt6-2* plants under the control of the constitutive 2×35S promoter. The accumulation of flavonol glycosides was analysed in herbizide-bleached seedlings using DPBA-staining (Figure 3B). While Col-0 wildtype seedlings appeared yellow under UV light, indicating the accumulation of flavonol glycosides, *tt6-2* seedlings showed a red fluorescence. *tt6-2* mutants expressing *MusaF3H1* or *MusaF3H2* were able to complement the mutant phenotype, showing the characteristic yellow flavonol glycoside fluorescence of the wild type. These results demonstrate that *Musa*F3H1 and *Musa*F3H2 are functional enzymes with F3H activity and are able to catalyse the conversion of naringenin to DHK.

**Figure 3:**
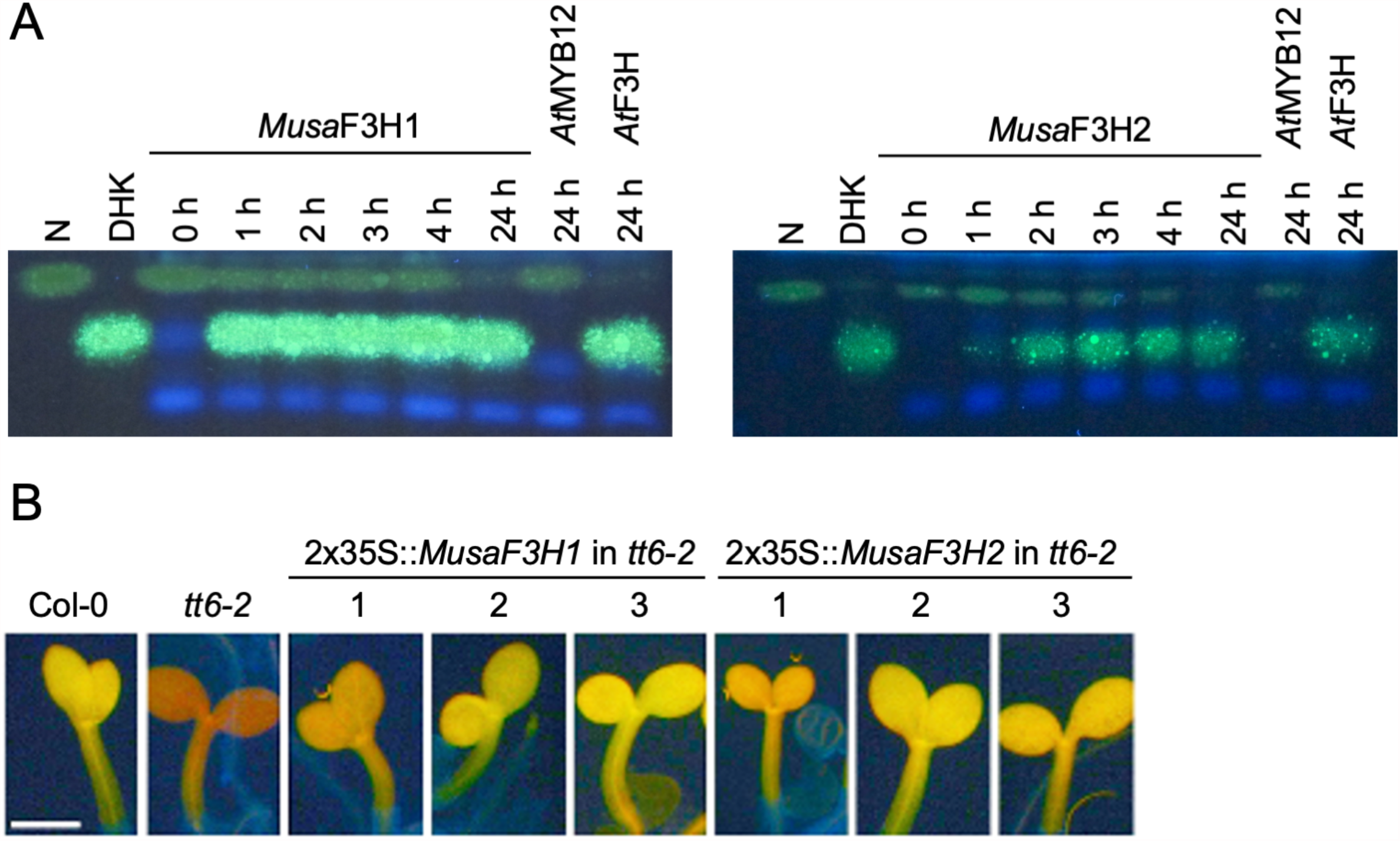
*Musa*F3H1 and *Musa*F3H2 are functional flavanone 3-hydroxylases. (A) HPTLC analysis of F3H bioconversion assays using extracts from *E. coli* expressing recombinant *Musa*F3H1 or *Musa*F3H2. The substrate naringenin (N) and the product dihydrokaempferol (DHK) were used as standards. *At*F3H was used as a positive control. *At*MYB12 was used as a negative control. (B) Analysis of *tt6-2* mutant seedlings complemented with *Musa*F3H1 and *Musa*F3H2. Col-0 wildtype and *tt6-2* were used as controls. Representative pictures of DPBA-stained seedlings under UV light are given. Yellow fluorescence indicates flavonol glycoside accumulation. The different numbers indicate individual transgenic lines. The scale bar indicates 0.5 mm.

### *Musa*FLS1 and *Musa*FLS3 are functional flavonol synthases

To confirm FLS activity *in vivo*, the enzymes were assayed in a coupled bioconversion experiment (Figure 4A). Two *E. coli* cultures expressing *MusaF3H1* and a *MusaFLS* were mixed and fed with naringenin in the presence of 2-oxoglutarate and ferrous irons. Here, *Musa*F3H1 converts naringenin to DHK, thus providing the substrate for the FLS enzyme to be tested. While *Musa*FLS1 was found to be able to convert DHK to kaempferol under the assay conditions, *Musa*FLS3 was not. In a second bioconversion experiment, *E. coli* cultures expressing *MusaFLS1* or *MusaFLS3* were fed with DHQ (Supplementary File S5). Again, *Musa*FLS1 was able convert the dihydroflavonol to the corresponding flavonol, while *Musa*FLS3 was not. We checked the expression of recombinant *Musa*FLS1 and *Musa*FLS3 in the *E. coli* cultures by SDS-PAGE and found both *Musa*FLS proteins being expressed at similar levels (Supplementary File S7). Moreover, we analysed a possible F3H/FLS biofunctional activity of *Musa*FLS1 and *Musa*FLS3 by feeding naringenin to *E. coli* cultures. While *Musa*FLS1 was not able to convert naringenin to DHK in the assay, *Musa*FLS3 showed F3H activity but not a further conversion to kaempferol (Supplementary File S6). To further analyse the enzymatic properties *in planta, MusaFLS1* or *MusaFLS3* were expressed in the flavonol and anthocyanin deficient *A. thaliana ans/fls1-2* double mutant. HPTLC analyses of methanolic extracts from rosette leaves from greenhouse-grown plants (Figure 4B) showed that wildtype plants (Col-0, Nö-0) contained flavonol glycosides, including the prominent derivatives kaempferol 3-O-rhamnoside-7-O-rhamnoside (K-3R-7R), kaempferol 3-O-glucoside-7-O-rhamnoside (K-3G-7R) and kaempferol 3[-O-rhamnosyl-glucoside[-7-O-rhamnoside (K-3[G-R]-7R). *ans/fls1-2* plants accumulated several dihydroflavonols but did not show flavonol derivatives. *ans/fls1-2* mutants transformed with 2×35S::*MusaFLS1* or 2×35S::*MusaFLS3* constructs were able to form several flavonol derivatives. Nevertheless, intensities and accumulation patterns of flavonol glycosides varied. We also analysed if *MusaFLS1* and *MusaFLS3* are able to complement the anthocyanin-deficiency of the *ans/fls1-2* double mutant. For this, seedlings were grown on 4 % sucrose to induce anthocyanin accumulation. As shown in Figure 4C, 6-day-old Col-0 and Nö-0 seedlings were able to accumulate red anthocyanin pigments, while *ans/fls1-2* transformed with *MusaFLS1* or *MusaFLS3* did not show visible anthocyanins. These results confirm that *Musa*FLS1 and *Musa*FLS3 are functional proteins with FLS activity and are enzymes able to catalyse the conversion of dihydroflavonol to flavonol.

**Figure 4:**
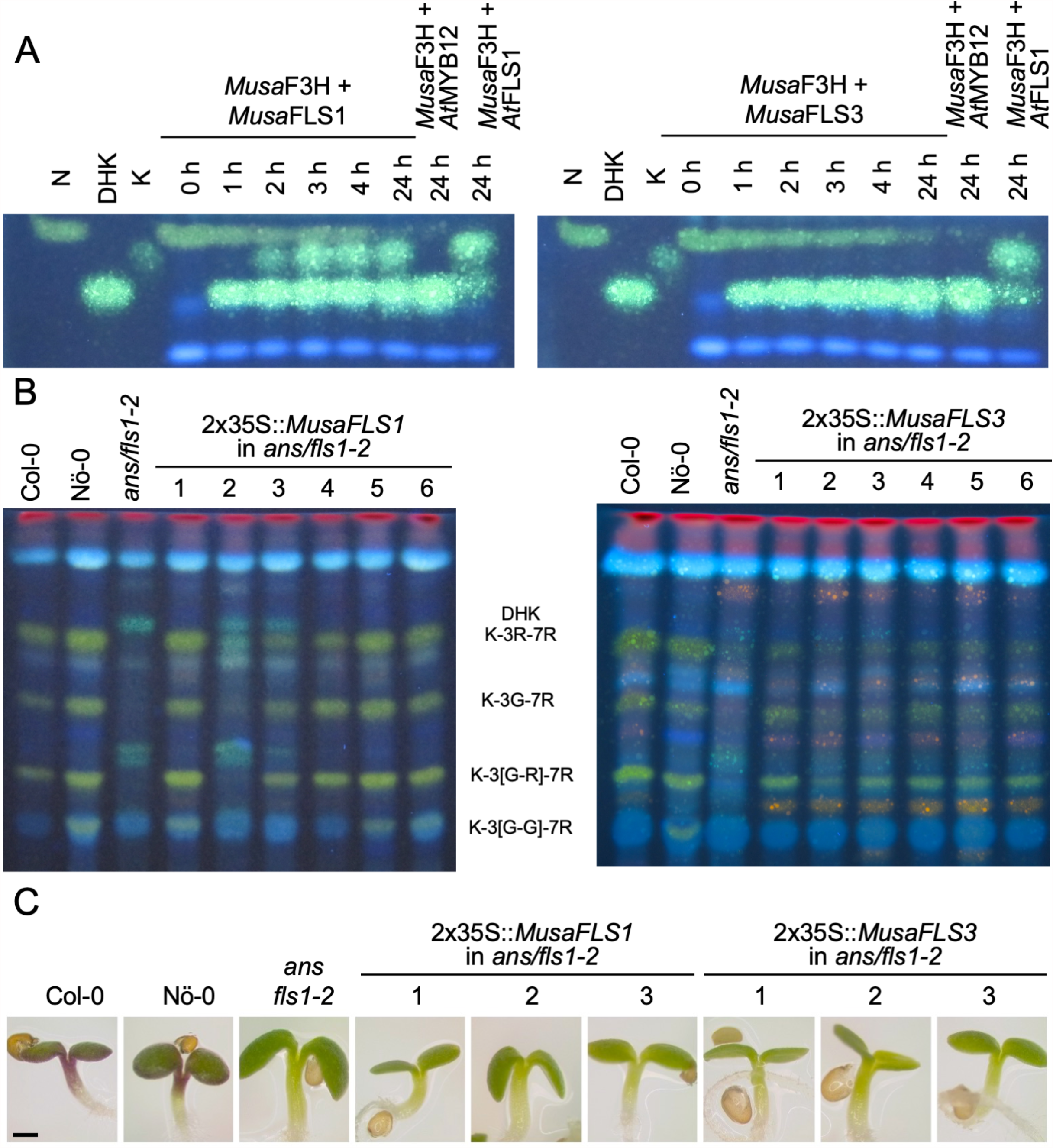
*Musa*FLS1 and *Musa*FLS3 are functional flavonol synthases. (A) HPTLC analysis of a FLS bioconversion assays using extracts from *E. coli* expressing recombinant *Musa*FLS1 or *Musa*FLS3. The F3H substrate naringenin (N), the FLS substrate dihydrokaempferol (DHK) and the product kaempferol (K) were used as standards. *At*FLS1 served as a positive control and *At*MYB12 was used as a negative control. (B) Flavonol glycoside accumulation in *Musa*FLS-complemented *ans/fls1-2* seedlings, analysed by HPTLC analysis. Col-0, Nö-0 (both wildtype) and *ans/fls1-2* were used as controls. Bright green spots belong to derivates of kaempferol, orange spots are derivates of quercetin and faint blue shows sinapate derivatives. Dark green and yellow spots indicate DHK and DHQ, respectively. G, glucose; K, kaempferol; Q, quercetin; R, rhamnose (C) Representative pictures of anthocyanin (red) accumulation in 6-day-old *Musa*FLS-complemented *ans/fls1-2* seedlings growing on 4 % sucrose. The scale bar indicates 0.5 mm.

### *Musa*ANS is a functional anthocyanidin synthase

We analysed *Musa*ANS functionality in a complementation assay with the *ans/fls1-2 A. thaliana* double mutant. To examine the ability of *MusaANS* to complement the *ans/fls1-2* anthocyanin deficiency phenotype, seedlings were grown on anthocyanin-inducing media. Anthocyanin accumulation was analysed visually and quantified photometrically (Figure 5A-B). While wildtype seedlings showed red pigmentation, the *ans/fls1-2* seedlings did not. *ans/fls1-2* seedlings expressing *MusaANS* showed accumulation of anthocyanins. The anthocyanin content in the complemented seedlings was strongly increased compared to *ans/fls1-2* knockout plants, indicating ANS activity. Furthermore, we analysed the ability of *MusaANS* to complement the flavonol deficiency phenotype of *ans/fls1-2* plants. For this, methanolic extracts of seedlings were analysed by HPTLC followed by DPBA-staining (Figure 5C). While wildtype seedlings accumulated several kaempferol and quercetin glycosides, *ans/fls1-2* mutants expressing *MusaANS* showed a flavonoid pattern identical to the *ans/fls1-2* mutant, accumulating dihydroflavonol derivatives, but no flavonol derivatives. These results indicate that *Musa*ANS is a functional enzyme with ANS activity and is able to catalyse the conversion of leucoanthocyanidin to anthocyanidin.

**Figure 5:**
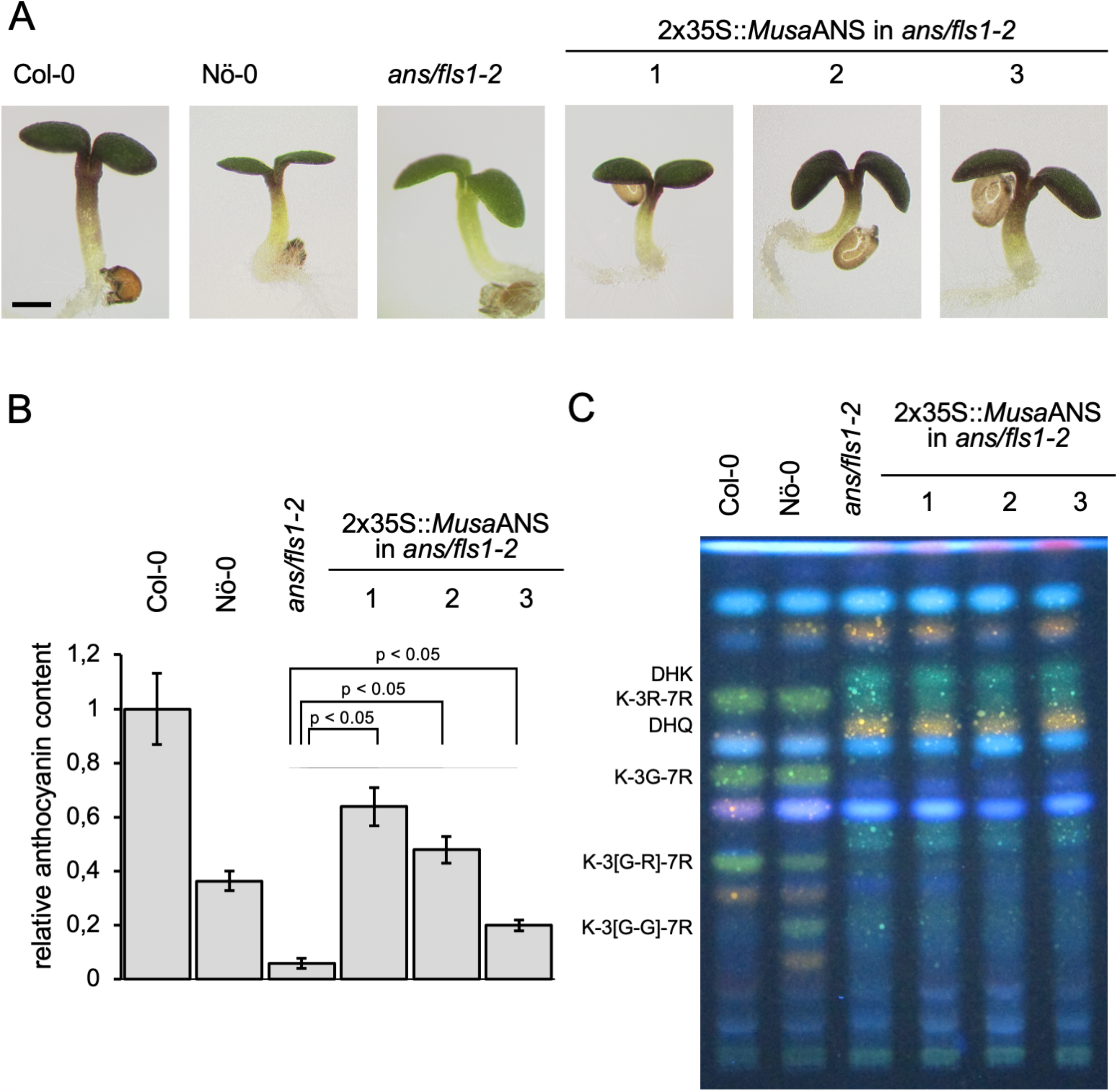
*Musa*ANS is a functional anthocyanin synthase. Analysis of *ans/fls1-2* double mutant seedlings complemented with 2×35S driven *MusaANS* demonstrate *in planta* ANS functionality of *Musa*ANS by anthocyanin accumulation. The different numbers indicate individual transgenic lines. (A-B) Sucrose induced anthocyanin accumulation in 6-day-old *A. thaliana* seedlings. (A) Representative pictures of seedlings (the scale bar indicates 0.5 mm) and (B) corresponding relative anthocyanin content. Error bars indicate the standard error for three independent measurements. (C) *Musa*ANS does not show *in planta* FLS activity. Flavonol glycoside accumulation in *MusaANS*-complemented *ans/fls1-2* seedlings, analysed by HPTLC analysis. Col-0, Nö-0 (both wildtype) and *ans/fls1-2* were used as controls. Bright green spots belong to derivates of kaempferol, orange spots are derivates of quercetin and faint blue shows sinapate derivatives. Dark green and yellow spots indicate DHK and DHQ, respectively. G, glucose; K, kaempferol; Q, quercetin; R, rhamnose.

## Discussion

In this study, we isolated several cDNAs of *in silico* annotated 2-ODD-type flavonoid biosynthesis enzyme coding genes from *Musa* and tested the *in vivo* functionality of the encoded proteins in *E. coli* and *A. thaliana*.

### *Musa*F3H1 and *Musa*F3H2

*In vivo E. coli* bioconversion assays revealed that *Musa*F3H1 and *Musa*F3H2 can covert naringenin to DHK. Moreover, *MusaF3H1* and *MusaF3H2* are able to complement the loss-of-function phenotype of *A. thaliana tt6-2* seedlings, showing *in planta* F3H-activity. Therefore, we conclude that *Musa*F3H1 and *Musa*F3H2 are functional flavanone 3-hydroxylases.

Previous studies annotated *Ma02_g04650* (*MusaF3H1*) and *Ma07_g17200* (*MusaF3H2*) as genes encoding flavanone 3-hydroxylases (Martin et al., 2016; Pandey et al., 2016; Pucker et al., 2020b). However, the automatic approach developed by Pucker et al. (2020b) also considered *Ma07_g17200* as a candidate to encode a FNSI enzyme. For a parsley F3H protein it has been found that a replacement of three amino acids was sufficient to cause FNSI side activity and seven amino acid exchanges almost lead to a complete change in enzyme activity towards FNSI functionality (Gebhardt et al., 2007). These findings underline the particular high similarity between FNSI and F3H. A closer inspection of the deduced peptide sequence of *Ma07_g17200* gene revealed a lack of conservation of amino acid residues known to be relevant for FNSI function (i.e. T106M, T115I, I116V, F131I, E195D, I200V, V215L, R216K). Furthermore, *Ma07_g17200* did not show FNSI activity in our bioconversion assay. Together with the confirmed F3H activity, this indicates that the classification of *Ma07_g17200* as a FNSI encoding gene was inaccurate. Even though Ma07_p17200 is no functional FNSI, flavone derivatives have been identified in *Musa* (Fu et al., 2018). In *Gerbera* and *Glycine max*, FNSII is responsible for the formation of flavones (Martens and Forkmann, 1999; Fliegmann et al., 2010). A candidate *Musa*FNSII, encoded by *Ma08_g26160*, has been identified (Pucker et al., 2020b) and is probably responsible for the accumulation of flavones in *Musa*. Since FNSII enzymes are NADPH-dependent cytochrome P450 monooxygenases (Jiang et al., 2016), the putative *Musa*FNSII (encoded by *Ma08_g26160*) was not studied in this work.

Our expression study that was based on published RNA-Seq data (Pucker et al., 2020a), revealed highest *MusaF3H1* transcript abundance in early developmental stages of pulp and intermediate stages of peel development, while highest *MusaF3H2* expression was found in adult leaves (Table 1). This is, at first glance, in contrast to the quantitative real-time PCR (qRT-PCR)-derived expression data presented by Pandey et al. (2016), reporting high transcript levels of *MusaF3H1* and *MusaF3H2* in young leaves and bract. In this study, the authors also show increased *MusaF3H1* expression in pseudostem, root and ripe pulp, compared to *MusaF3H2*. These discrepancies are probably due to the different growth conditions, germplasms/cultivars and sampling time points in the generation of the expression data and cannot be reasonably analysed further at this point. It should be noted, however, that *MusaF3H2* expression in leaves of plantlets was found to be increased following treatment with the phytohormone methyl jasmonate (MJ), while *MusaF3H1* expression was not (Pandey et al., 2016). As MJ is involved in various regulatory processes, including the response against biotic and abiotic stresses (summarised in Cheong and Choi, 2003), the MJ-dependent induction of *MusaF3H2* expression could imply that the *Musa*F3H2 enzyme plays a specialised role in the formation of flavonoids in response to stresses. Such stress response induction of F3H encoding genes was previously shown for a *F3H* gene from the dessert plant *Reaumuria soongorica*, which is induced by UV-light (Liu et al., 2013) and two *F3H* genes from *Camellia sinensis* (*C. sinensis*), which are induced by UV light and by treatment with abscisic acid (ABA) or sucrose (Han et al., 2017). In addition, overexpression of *F3H* from *C. sinensis* (Mahajan and Yadav, 2014) and from *Lycium chinense* (Song et al., 2016) in tobacco improved the tolerance to salt stress and fungal pathogens in the first case and to drought stress in the latter case. In conclusion, our results clearly show that *Musa*F3H1 and *Musa*F3H2 are functional flavanone 3-hydroxylases and that *Musa*F3H2 might play a role in stress response.

### *Musa*FLS1 and *Musa*FLS3

*Musa*FLS1 was found to be able to convert DHK and DHQ to the corresponding flavonol in the *E. coli* bioconversion assays. This ability was validated *in planta* by successful complementation of the flavonol deficiency of *A. thaliana ans/fls1-2* double mutant seedlings, while the anthocyanin deficiency phenotype was not restored. These observations could hint to an exclusive FLS activity of *Musa*FLS1. FLS and DFR, the first enzymes of the flavonol and anthocyanin branches of flavonoid biosynthesis, compete for dihydroflavonol substrates (Figure 1). This is demonstrated by several *A. thaliana fls1* single mutants which accumulate higher levels of anthocyanin pigments (Owens et al., 2008; Stracke et al., 2009). Falcone Ferreyra et al. (2010) found that overexpression of *ZmFLS1* in an *A. thaliana fls1* mutant decreases the accumulation of anthocyanins to wildtype level, again indicating that the anthocyanin and flavonol branches of flavonoid biosynthesis can compete for dihydroflavonol substrates in the same cell or tissue. Accordingly, the overlapping substrate usage of FLS and ANS is difficult to analyse in a *fls1* single mutant. Here, the use of an *A. thaliana ans/fls1-2* double mutant is a simple way to analyse the FLS and a possible ANS side activity of flavonol synthases and vice versa *in planta*. To analyse further side activities, a *f3h fls ans* mutant would be beneficial. *A. thaliana ans/fls1-2* plants expressing *MusaFLS3* showed the accumulation of several kaempferol derivatives. In contrast, plants complemented with *MusaFLS1* also accumulate quercetin derivatives. This is most likely an effect of different environmental conditions in plant growth. One set of plants was grown in the greenhouse (*MusaFLS1*), another set was grown in a growth chamber (*MusaFLS3*). In the plants used for *Musa*FLS3 experiments these conditions promoted the accumulation of DHQ, while the other growth conditions did not in the plants used for *MusaFLS1* experiments. Accordingly, DHQ was probably not available as a substrate for *Musa*FLS1 and could not be converted to quercetin. A possible explanation for the varying DHQ amounts could be the light induced expression of F3’H, which converts DHK to DHQ. The expression of *F3’H* in cultured *A. thaliana* cells can be induced by UV-light (Schoenbohm et al., 2000) and the activity of the *F3’H* promoter from *Vitis vinifera* is increased under light exposure (Sun et al., 2015).

Consequently, a different light quality or higher light intensity could be responsible for increased *F3’H* expression, causing the accumulation of DHQ in the plants grown in the growth chamber. While *MusaFLS3* was able to complement the flavonol deficiency in the *ans/fls1-2* mutant, it did not lead to an accumulation of anthocyanins. In contrast to the *in planta* results, *Musa*FLS3 did not convert DHK or DHQ to the corresponding flavonol in the *in vivo* bioconversion assays, even though the protein was successfully expressed in the *E. coli* culture. We used the bioconversion assays as a simple and versatile tool to analyse enzymatic activities. Nevertheless, heterologous expression of eucaryotic proteins in *E. coli* is an artificial system. It can cause a divergent pattern of posttranslational modifications or production of high amounts of protein in inclusion bodies, leading to inactive protein (Sahdev et al., 2008). Preuss et al. (2009) reported that *At*FLS3 did not show FLS activity in *E. coli*, but did convert dihydroflavonols to the corresponding flavonols upon expression in yeast, indicating that the stability of *At*FLS3 was improved under the latter assay conditions. The approach carried out in this work might have similar limitations. Further limitations can be caused by different substrate preferences, as reported for *Os*FLS (Park et al., 2019). Despite the lack of FLS activity in the bioconversion assay, *Musa*FLS3 showed F3H activity. Nevertheless, the *in planta* complementation of the flavonol deficiency in the *A. thaliana ans/fls1-2* mutant and the lacking complementation of the anthocyanin deficiency phenotype in the *ans/fls1-2* mutant indicate that *Musa*FLS3 is a functional flavonol synthase with F3H side activity, which does not exhibit significant ANS activity.

*MusaFLS1* transcript abundance was found to be high in adult leaves and low in roots. *MusaFLS3* and *MusaFLS4* revealed opposed transcript levels. Such divergent expression patterns have previously been observed for *FLS1* and *FLS2* from *Freesia hybrida* and *FLS1* and *FLS2* from *Cyclamen purpurascens* (Akita et al., 2018; Shan et al., 2020). These opposed expression patterns could point to differential activity in distinct organs. Furthermore, the tandemly arranged *MusaFLS3* and *MusaFLS4* genes show very similar expression patterns (Table 1), possibly indicating functional redundancy. While we could not find expression of *MusaFLS2* in our expression analyses, Pandey et al. (2016) observed *MusaFLS2* expression in several organs (including bract, pseudostem and root), as well as in different developmental stages of peel and pulp. Again, these results show that different cultivars, growth conditions, sampling time points and analysed organs can have a strong influence on the resulting data. Accordingly, data from different studies should be evaluated carefully. *MusaFLS1* and *MusaFLS2* expression in leaves of plantlets does not significantly increase after MJ treatment (Pandey et al., 2016). However, UV radiation induces the expression of FLS from *Z. mays* (Falcone Ferreyra et al., 2010) and *M. domestica* (Henry-Kirk et al., 2018) and the relative expression of FLS from *Triticum aestivum* increases during drought stress (Ma et al., 2014). Together with the knowledge that flavonols act as antioxidants (Wang et al., 2006), an involvement of *Musa*FLSs in stress response seems feasible and should be further analysed under a broader range of conditions. In summary, our results show that *Musa*FLS1 and *Musa*FLS3 are functional flavonol synthase enzymes and hint at possible differential organ specific activities of *Musa*FLS1 and *Musa*FLS3 / *Musa*FLS4.

### *Musa*ANS

*A. thaliana ans/fls1-2* seedlings expressing *MusaANS* show a strong, red pigmentation, revealing that *MusaANS* can complement the anthocyanin deficiency caused by mutation of *AtANS*. However, the seedlings did not display flavonol derivatives in HPTLC analyses, which have been reported to detect flavonol glycosides at levels of 50 pMol (Stracke, 2010). These results indicate that *Musa*ANS is a functional anthocyanidin synthase, but does not have FLS activity. The RNA-Seq data-derived expression profiles revealed high *MusaANS* transcript abundance in pulp and peel. Additionally, *MusaANS* expression has been reported to be high in bract tissue (Pandey et al., 2016), the specialised leaves surrounding the flowers and usually coloured red or purple due to anthocyanin accumulation (Pazmiño-Durána et al., 2001). This spatial correlation of *MusaANS* transcripts and anthocyanin metabolites in bract tissue supports the proposed biological functionality of this enzyme, catalysing the conversion of leucoanthocyanidins to anthocyanidins in *Musa. MusaANS* expression increases 24 hours after MJ treatment and decreases following dark treatment (Pandey et al., 2016). As supposed for *MusaF3H2*, the increased expression of *MusaANS* after MJ treatment could imply an involvement in the formation of anthocyanins as a consequence of stress response. ABA, salicylic acid (SA), UV-B and cold treatments have been shown to enhance *ANS* transcript abundance in *G. biloba* (Xu et al., 2008) and overexpression of *OsANS* raises the antioxidant potential in *O. sativa* (Reddy et al., 2007). The accumulation of anthocyanins has often been shown to be induced by (UV-) light (Takahashi et al., 1991; Stapleton and Walbot, 1994; Meng et al., 2004). We therefore assume that also *Musa*ANS could be involved in such stress response.

Very recently, a R2R3-MYB-type transcription factor (MaMYB4) has been identified as a negative regulator of anthocyanin biosynthesis in *Musa* acting as repressor on the *MusaANS* promoter (Deng et al., 2021). These results give a first insight into the transcriptional regulation of *MusaANS* expression and confirm the role of *MusaANS* in the anthocyanin biosynthesis in *Musa*. Taking all available evidence into account, *MusaANS* encodes a functional anthocyanidin synthase with a possible involvement in the plants’ stress response.

To deepen the knowledge about 2-ODDs and other enzymes involved in *Musa* flavonoid biosynthesis, it would be beneficial to acquire even more spatially and timely highly resolved transcriptome and in particular metabolite data. This data could serve as a starting point for the analysis of organ-, stress- or development-specific enzyme activities and functions, as well as possible substrate preferences. It could also be used to elucidate the regulatory network of *Musa* flavonoid biosynthesis. Furthermore, knowledge about the influence of different stresses (e.g., pathogens, light, temperature) on specific transcriptomes and metabolomes of the *Musa* plant could help to widen the knowledge of flavonoid biosynthesis and particular the functionality of the *Musa* 2-ODD enzymes. This could also lead to the detection of possible restricted side activities or overlapping functionalities as described for 2-OODs in some other plant species (Falcone Ferreyra et al., 2010; Park et al., 2019) and to further decode the cause of these multifunctionalities in 2-ODDs.

To conclude, in this study the functionality of five 2-ODDs involved in flavonoid biosynthesis in *Musa* was demonstrated *in vivo* in bacterial cells and *in planta*. Knowledge gained about the structural genes *MusaF3H1, MusaF3H2, MusaFLS1, MusaFLS3* and *MusaANS* in a major crop plant provides a basis for further research towards engineered, increased flavonoid production in banana, which could contribute to the fruits’ antioxidant activity and nutritional value, and possibly even enhance the plants’ defence against *Foc*-TR4.

## Author Contributions

MB and RS planned the experiments. MB and CA performed the experiments and analysed the data. MB and SM interpreted the TLC data. RS and BW supervised the project. MB wrote the initial draft. RS and BW revised the manuscript. All authors read and approved the final manuscript version.

## Acknowledgements

We are grateful to Melanie Kuhlmann for her excellent assistance in the laboratory and to Andrea Voigt for her competent help in the greenhouse. We thank Anika Beckers who contributed to creation of constructs with *MusaFLS* cDNAs and Prisca Viehöver for sequencing. In addition, we thank Thomas Baier for his support with the SDS-PAGE.

## Supplementary Material

Table S1: Plant 2-ODDs with proven F3H, FNSI, FLS and ANS functionality, used in phylogenetic analysis.

Table S2: Sequence Read Archive IDs for RNA-Seq raw data used for expression analysis.

Table S3: Oligonucleotide primers, used in this work.

File S1: CDS of *Musa2-*ODDs from this work.

File S2: Multiple sequence alignment of flavonoid biosynthesis related 2-ODDs from banana and plant proteins with demonstrated 2-ODD activity.

File S3: *Musa*F3H1 and *Musa*F3H2 convert eriodictyol to dihydroquercetin.

File S4: Analysis of FNSI side activity of *Musa*F3H2.

File S5: MusaFLS1 converts dihydroquercetin to quercetin.

File S6: Analysis of F3H side activity of *Musa*FLS1 and *Musa*FLS3.

File S7: Analysis of recombinant *Musa*FLS1 and *Musa*FLS3 in *E. coli* by SDS-PAGE.

