## Supplementary for "Functional characterisation of banana (*Musa* spp.) 2-oxoglutarate-dependent dioxygenases involved in flavonoid biosynthesis"

**Supplementary material**

**Table S1: Plant 2-ODDs with proven F3H, FNS I, FLS and ANS functionality, used in phylogenetic analysis.**

| **Name** | **Species** | **GenBank ID** | **Reference** | **Function** |
| --- | --- | --- | --- | --- |
| MtF3H | *Medicago truncatula* | FJ529406.1 | Shen et al. (2010) | F3H |
| OsF3H | *Oryza sativa* | NM_001060692.1 | Kim et al. (2008) | F3H |
| PcF3H | *Petroselinum crispum* | AY230248.1 | Martens et al. (2001) | F3H |
| PhF3H | *Petunia hybrida* | AF022142.1 | Britsch et al. (1986) | F3H |
| ZmF3H | *Zea mays* | AAA91227.1 | Deboo et al. (1995) | F3H |
| AcFLS | *Allium cepa* | KY369209.1 | Park et al. (2017) | FLS |
| AtFLS | *Arabidopsis thaliana* | NP_001190266.1 | Owens et al. (2008) | FLS |
| CuFLS | *Citrus unshiu* | Q9ZWQ9.1 | Wellmann et al. (2002) | FLS |
| FaFLS | *Fragaria x ananassa* | DQ087252 | Almeida et al. (2007) | FLS |
| MdFLS | *Malus domestica* | AY965343 | Halbwirth et al. (2006) | FLS |
| OsFLS | *Oryza sativa* | XP_015624815.1 | Park et al. (2019) | FLS |
| PcFLS | *Petroselinum crispum* | Q7XZQ6.1 | Martens et al. (2003) | FLS |
| PhFLS | *Petunia hybrida* | Q07512.1 | Holton et al. (1993) | FLS |
| ZmFLS | *Zea mays* | BT039956 | Falcone Ferreyra et al. (2010) | FLS |
| AtDMR6 | *Arabidopsis thaliana* | OAO94377.1 | Falcone Ferreyra et al. (2015) | FNSI |
| CjFNSI | *Conocephalum japonicum* | QEP99662.1 | Li et al. (2020) | FNSI |
| CmFNSI | *Conium maculatum* | AY817677 | Gebhardt et al. (2005) | FNSI |
| MeFNSI | *Marchantia emarginata* | QEP99659.1 | Li et al. (2020) | FNSI |
| MpaFNSI | *Marchantia paleacea* | QEP99658.1 | Li et al. (2020) | FNSI |
| PaFNSI | *Plagiochasma appendiculatum* | QEP99657.1 | Li et al. (2020) | FNSI |
| PcFNSI | *Petroselinum crispum* | Q7XZQ8.1 | Martens et al. (2001) | FNSI |
| ZmFNSI | *Zea mays* | NP_001151167.1 | Falcone Ferreyra et al. (2015) | FNSI |
| AcANS | *Allium cepa* | EF192598.1 | Kim et al. (2004) | ANS |
| AtANS | *Arabidopsis thaliana* | NP_194019.1 | Wilmouth et al. (2002) | ANS |
| FaANS | *Fragaria x ananassa* | AY695817.1 | Almeida et al. (2007) | ANS |
| GbANS | *Ginkgo biloba* | ACC66092.1 | Xu et al. (2008) | ANS |
| OsANS | *Oryza sativa* | Y07955.1 | Reddy et al. (2007) | ANS |
| PaANS | *Phytolacca americana* | AB198870.1 | Shimada et al. (2005) | ANS |
| PfANS | *Perilla frutescens* | O04274.1 | Saito et al. (1999) | ANS |
| PhANS | *Petunia hybrida* | AB027454.1 | Nakajima et al. (2001) | ANS |
| SoANS | *Spinacia oleracea* | AB198869.1 | Shimada et al. (2005) | ANS |
| TfANS | *Torenia fournieri* | AB044091.1 | Nakajima et al. (2001) | ANS |
| ZmANS | *Zea mays* | NP_001106074 | Nakajima et al. (2001) | ANS |
| AtGA3ox1 | *Arabidopsis thaliana* | NP_173008.1 | Kawai et al. (2014) | outgroup |

**Table S2: Sequence Read Archive IDs for RNA-Seq raw data used for expression analysis.**

| **biolocial source** | **Sequence Read Archive IDs** |
| --- | --- |
| embryogenic | SRR8761343, SRR8761344, SRR8761345 |
| seedling | SRR7340404, SRR7340406 |
| root | SRR2984591, SRR2984594, SRR2984596, SRR2984598, SRR2984601, SRR4450942, SRR4450934, SRR4450941, SRR4450933, SRR4450944 |
| leaf | SRR6320573, SRR6320439, SRR6320438, SRR6320441, SRR6320440, SRR6320443, SRR5405140, SRR5405141 |
| young_leaf | SRR6320573, SRR6320439 |
| adult_leaf | SRR6320438, SRR6320441 |
| old_leaf | SRR6320440,SRR6320443 |
| pulp | SRR5405133, SRR6320444, SRR5405132,SRR6320447, SRR5405131, SRR6320446, SRR5405130, SRR6320531 |
| pulp S1 | SRR5405133, SRR6320444 |
| pulp S2 | SRR5405132, SRR6320447 |
| pulp S3 | SRR5405131, SRR6320446 |
| pulp S4 | SRR5405130, SRR6320531 |
| peel | SRR5405139, SRR5405138, SRR5405137, SRR6320442, SRR5405136, SRR6320445, SRR5405135, SRR5405134 |
| peel S1 | SRR5405139, SRR5405138 |
| peel S2 | SRR5405137, SRR6320442 |
| peel S3 | SRR5405136, SRR6320445 |
| peel S4 | SRR5405135, SRR5405134 |

**Table S3: Oligonucleotide primers, used in this work.**

| **Name** | **Sequence (5’-3’)** | **Description** |
| --- | --- | --- |
| RS1708 | attB1-ccATGGCTCCCGGCACCGCAGTCCTTCCC | fwd Primer with attB1 for *Musa*F3H1 (Ma02_t04650) CDS |
| RS1709 | *attB2-*aAGCCAAGATCTCATCGAGGCCTGTAGCC | rev Primer with attB2 for *Musa*F3H1 (Ma02_t04650) CDS |
| RS1710 | *attB1-*ccATGGCCCCCGGCACTGCATTGCTTCCC | fwd Primer with attB1 for *Musa*F3H2 (Ma07_t17200) CDS |
| RS1712 | *attB2-*aAGCTAGGATCTCATTTAGGCATTTCGCC | rev Primer with attB2 for *Musa*F3H2 (Ma07_t17200) CDS |
| RS1681 | *attB1-*ccATGGAGGTGGAGAGGGTGCAGGCC | fwd Primer with attB1 for *Musa*FLS1 (Ma03_t06970) CDS |
| RS1682 | attB2-aCTGCGGAAGCTTGTTCAGCTTGCAG | rev Primer with attB2 for *Musa*FLS1 (Ma03_t06970) CDS |
| RS1695 | attB1-ccATGGAGGTGGAGAGGGTTCAAGTCATCGCG | fwd Primer with attB1 for *Musa*FLS3 (Ma10_t25110) CDS |
| RS1696 | attB2-aTTGCGGAAGCTTGTTGAGCTTGCAGTAGGCG | rev Primer with attB2 for *Musa*FLS3 (Ma10_t25110) CDS |
| RS1704 | attB1-ccATGGCCACCAAGGTCGTGTCTG | fwd Primer with attB1 for *Musa*ANS (Ma05_t03850) CDS |
| RS1705 | attB2-aGTTTGGGGTCTTGAAGTCCCCCCG | rev Primer with attB2 for *Musa*ANS (Ma05_t03850) CDS |
| RS2101 | attB1-ccATGGCTCCTACAACAATAAC | fwd Primer with attB1 for *PcFNSI* CDS |
| RS2102 | attB2-AGCTAAATTTTCATCTGCAC | rev Primer with attB2 for *PcFNSI* CDS |

attB1: GGGGACAAGTTTGTACAAAAAAGCAGGCT, attB2: GGGGACCACTTTGTACAAGAAAGCTGGGT

**File S1: CDS of *Musa2-*ODDs from this work.**

Variant positions to the Pahang reference genome are given in bold-underlined.

*Musa*F3H1

ATGGCTCCCGGCACCGCAGTCCTTCCCACCAACGAGCAGACCTTGAGGGCGAGCTTCGTCCGGGACGAGGACGAGCGTCCCAAGGTGGCTTACAACCAGTTCAGCCTTGACATCCCCGTGATCTCGCTCTCCGGGATTGATGACGACGCCGCGGGCGGGAAGAGGGCGGAGATCCGCCGCAAGGTCGTGGAGGCCTGCGAGGACTGGGGTATATTCCAGGTGGTGGACCACGGGGT**G**GACGCCGGCCTTATCTCCGACATGACTAGGCTGGCCAAAGAGTTCTTCGCGCTGTCCCCGGA**T**GAGAAGCTCTTGTTCGACATGTCCGGTGGGAAGAAGGGAGGGTTCATCGTCTCCAGCCACCTCCAGGGCGAGGCAGTTCAGGACTGGAGAGAGATCGTGACGTACTTCTCATACCCAATTCGGGCTCGGGACTACTCGCG**G**TGGCCGGACAAGCCGGAAGGGTGGAGATCCGTGGTGGAGTCGTACAGCGAGAAGCT**G**ATGGGCTTGGCGTGCAAGCTCTTGGAGGTGTTGTCTGAGGCTATGGGGCTCGACAAGGAGGCCCTCACCGAGGCCTGCATCGACATGGATCAGAAGGT**G**GTCGTCAACTACTACCCCAAGTGCCCCCAGCCGGACCTCACTCTTGGCCTCAAACGACACACCGATCCCGGCACCATCACCCTCCTCCTCCAAGACCAAGTCGGCGGCCTCCAAGCCACCAAGGATGGCGGCAAGACATGGATCACGGTTCAGCCTGTGGAGGGAGCTTTTGTGGTCAACCTCGGGGACCATGGCCATTTTCTGAGCAATGGGAGGTTCAAGAATGC**A**GACCACCAGGCTGTGGTGAACTCAAACT**G**CAGCAGGCTCTCCATCGCCACGTTTCAGAACCCCGCGCCGGAGGCAATCGTGTATCCGCTGGCAGTAAGGGAGGGAGAGAAGCCGATTCTGGA**C**GAGCCCATCACGTTTGCCGAGATGTACCGCAGGAAGATGAGCCGAGACCTCGAGCTCGCCAAGCTCAAGAAACTGGCCAAGACGGAGCAG**C**AGCAG**A**AGCCGGAGCTACT**A**GAGAAGACTAAGGACATCAACTTGGCCAAGGCTACAGGCCTCGATGAGATCTTGGCT

*Musa*F3H2

ATGGCCCCCGGCACTGCATTGCTTCCCACGAACGAGGCGACCCTGAGGGCGAGCTTCGTGCGGGACGAGGACGAGCGACCCAAGGTGGCCTA**C**AACCA**G**TTCAGCAGCGACATCCC**C**GTGATCTCGCTCGCCGGGGTTGACGACGAGGACGGGGCCACCGGCGGGCGGAGGGCGGAGATCCGCCGCAAGATCGTGGA**A**GCCTGCGAGGACTGGGGCTTATTCCAGGTG**T**TGGACCATGGAGTGGACGCCGGCATGATCTCCGAGATGACCCGGCTCGCCCGGGATTTCTTCGCGCTACCCCCCGAGGAGAAGCT**A**CGGTTCGACATGTCCGGCGGAAAAAAGGGAGGCTTCATCGTGTCCAGCCATCTCCAGGGTGAGGCAGTGCAGGACTGGCGAGAGATCGTGACTTACTTCTCGTACCCGATCGGAGCTCGGGACTACTCGAGGTGGCCGGATAAGCCAGAAGGGTGGAGGTCGGTGGTGGAGTCATACAGCGAGAAGCTGATGGGGTTGGCGTGCAAGCTGCTGGAGGTGTTGTCGGAGGCAATGGGGCTCGACAAGGAAGCCCTCAC**T**GATGCCTGCGTAGACATGGATCAGAAGGTGGTGGTCAACTACTACCCCA**T**GTGCCCCCAGCCCGACCTCACCCTCGGCCTCAAACGCCACACCGATCCCGG**C**ACCATCACCCTCCTCCTCCAAGACCAAGTCGGCGGACTCCAAGCCACCAAGGACGGCGGCAAGACGTGGATCACAGTTCAGCCCGTGGAGGGAGCTTTCGT**T**GTCAACCTCGGGGACCATGGCCATTATCTGAGCAACGGGAGGTTCAAGAACGC**C**GATCATCAGGCGGTGGTGAACTCGAGCTGCAGCAGGCTGTCGATCGCCACGTTTCAGAACCCAGCGCCGGAGGCGATCGTGTATCCCCTGGCGATAAGGGAGGGAGA**G**AAGCCGATCCTGGACGAGCCCATCACGTTCAGCGAGATGTACCGTAGGAAGATGAGCCGAGATCTC**A**AGCTTGCCAACCT**T**AAGAA**G**CTGGCCA**A**GGCGGAGCATCAGCGGC

*Musa*FLS1

ATGGAGGTGGAGAGGGTGCAGGCCATCGCGTCCCTGAGCGTGGCCACCAACGACATACCGCCGGAGTTCGTGAGGTCGGAGCACGAGCAGCCGGGTATCACCACGTACCGCGGCC**T**GGTCCCGGAGATCCCGGTGATCGACCTCGAAGACGGGGACGAAGGCCGGGTGACGCGCGCCATCGCGGAGGCCAGCCAGGAGTGGGGCATCTTCCAGCTGGTGAACCACGGCATCCCCGGGGAGGTGATCCGGGCGCTGCAGCGCGTGGGCAGGGAGTTCTTCGAGCTGCCACCGGAGGAGAAGGAGAAGTACGCGGCGGCGCCGGGGAGCCTCCAGGGCTACGGAACCAAGCTGCAGAAGGACTTGGAAGGCAAGAAGGCGTGGGTGGACTTCCTCTTCCACAACATCTGGCCGCCGACGCACGTCGACCACCGCGCATGGCCGGAGAATCC**G**GTGGATTACAGGAAGGCAAATGAGGAGTACGCCAAACATTTGGTGGGATTGGTGGAGAAGATGTTGGTAAGCCTGTCCAAGGGACTGGGGCTGGAGGCCGACGTCCTCAAGCACGCAGTGGGAGGGGACGACTTGGAGTTCCTCCTCAAGATCAACTACTACCCGCCGTGCCCGAGACCCGACCTCGCCCTCGGCGTGGTGGCTCACACCGACATGTCCGCCATCACCATCCTGATCCCCAACGACGTCCCCGGCCTCCAGGTCTTCAAGGACGACCACTGGTTCGACGCCAAGTACGTCCCCGACGCCATCATCGTCCACATCGGGGACCAGATCGAGAAACTGAGCAACGGCAGGTACAAGAGCGTGCTGCACCGGACGACGGTGAACAAGGAGAAGGCGAGGATGTCGTGGCCTGTGTTCTGCTCCCCGCCGGGCGAGACGGTCATTGGGCCTCTGCCGCAGCTCGTCAGCGACGAACAGCCCGCTCAGTACAAGACGAAGAAGTACAAGGACTATGCTTTCTGCAAGCTGAACAAGCTTCCGCAG

*Musa*FLS3

ATGGAGGTGGAGAGGGTTCAAGTCATCGCTTCCGTCTGCGCCGCCGACGGTGTTATGCCTCCGGAGTTCATACGGTCGGAGCACGAGCAGCCGGGCATCACCACCTACCGCGGCCCCGCGCCGGAGATCCCCGTCATCGACCTCGCGGGCGCCGACCGGGA**C**CGGTTGACGATCGCCGTCGCCGAGGCCAGCCGGGAGTGGGGAATCTTCCAACTGCTGAACCACGGGATCCCGAGGGAGGTGATCCGTGAGCT**T**CAGCGCGTCGGCAAG**C**AATTCTTCGAGTTGCCGCAGGAGGAGAAGGAGATGTACGCGATGGAAT**G**CAAACCGGGGAGCTCGGAAGGATATGGGAC**C**AAGCTGCAGAGGGAGTTGGAGGGCAAAAAGGC**T**TGGGT**A**GACTTCTTCTTCCACTAC**A**T**T**TCGCCGCCGGCTCGCGTCAACCACGCCATCTGGCCCAAGAACCCT**G**CTGATTA**T**AGGAAAGCAAACGAGGAATATG**G**CAAACACCTGGTGGG**T**CTGGTGGACAAGATGCTGA**T**GAC**G**CTGTCGAGGGGACTGGGACTGGAGGAGCATGTCCTCAAGGGAGCACTCGGTGGAGATGGACTGGAGCTACG**T**CTAAAGATGAACTACTACCCACCGTGTCCTCGGCCTGACCTGGCTCTCGGCGTGGTGGCGCACACTGACATGTGTGCTATCACCTTCCTCGTCCCTAACCTTGTGCCGGGTTTGCAGGTCTTCAAAGATGAGCACTGGATTGACGTCAACTTCATTCCTAATGCTGTCATCGTCCACATCGGTGATCAGATCGAGATTTTGAGCAATGGAACATACAAGAGCGTGCTGCACAGAACGACCGTGAACAAAGAGAAGGTGAGGATGTCTTGGCCAGTGTTCTGCGCGCCTCCTGGTGAGATGGTAATTGGCCCTCTGCAACAGCTCGTCGGCGATGAGAGCCCAGCCAAGTATAAGCCGAAGAAGTACAAAGACTACGCCTACTGCAAGCTCAACAAGCTTCCGCAA

*Musa*ANS

ATGGCCACCAAGGTCGTGTCTGTGGCGCCCAGGGTGGAGATCCT**G**GC**G**AAGAGCGGCATCAACGAGATCCCGACCGAGTACGTCCGCCCCGAGTCGGAGCGGCTCGACCTCGGTGACGCGTTCGAGGAGGTGAAGAAGGCGGCGGAGGGGCCTCAGATTCCTGTGGTGGACCTCCAGGGTTTCGACTCGCCTGATGAAGAGGTGAGAAGGGCGTGCGTGGAGGAGGTGAGGAAGGCGGCGACGGAGTGGGGGGTGATGCACATCGTGAATCACGGCATCCCGTTGGAGCTCGTCGAGCAGCTGAGGAAGGTGGGGAAGGAGTTCTTCGACCTGCCCATAGAGCAGAAGGAGCAGTACGC**A**AACGACCAGTCATCAGGGAAGATCCAAGGGTACGGGAGCAAGCTGGCGAACAACGCGAGTGGGCAGCTCGAGTGGGAGGACTACTTCTTCCACCTTATATTCCCGGAGGAGAAGATCAACATCTCCATTTGGCCCAAGCAACCAACCGACTACATTGAGGTGACGAAGAAGTTCGGGAGGCAGCTGAGGGCGGTGGTCACCAAGATGCTGGAAGTTCTTTCCCTGGGTCTTGGACTGGAAGAGGGGAAGCTGGACAGGGAACTCGGAGGGATGGAGGACCTGCTGATGCAGTTGAAGATCAACTACTACCCTATCTGCCCACAGCCCGATCTCGCCCTCGGCGTCGAGGCGCACACCGACATCAGCGCGCTCTCCTTCATCCTCCACAACATGGTGCCGGGGCTGCAGATCTACTACGGCGGCAGGTGGGTCACCGCCAAATG**T**GTGCCGGACTCCATCATCATGCACGTCGGAGACTGCCTCGAGATCCTGAGCAATGGGCAGTACAAGAGCATCCTCCACCGCGGGCTCGTCAACAAGGAGAAGGTGCGCATCTCCTGGGCGGTCTTCTGCGAGCCTCCCAAAGACAAGATCGTGCTGAAGCCACTGGAGGAGCTCGTGGCCGACGGGACGCCGGCCAAGTTCCCCCCGCGCACCTTCGAGCAGCACATCCAGCACAAGCTCTTCAAGAAGACTCGGGGGGACTTCAAGACCCCAAAC

**File S2: Multiple sequence alignment of flavonoid biosynthesis related 2-ODDs from banana and plant proteins with demonstrated 2-ODD activity.**

Substrate binding residues are highlighted in grey. Bold-underlined amino acids are involved in binding ferrous iron or 2-oxoglutarate (Chua et al., 2008). Asterisks mark conserved residues, colons indicate very similar amino acids and full stops indicate weakly similar properties.

```
AtF3H           -------MAP-----GTLTELA---GESKLNSKFVRDEDERPKVAYNV----------FS	35
MusaF3H1        -------MAP-----GTAVLPT---NEQTLRASFVRDEDERPKVAYNQ----------FS	35
MusaF3H2        -------MAP-----GTALLPT---NEATLRASFVRDEDERPKVAYNQ----------FS	35
OsF3H           -------MAPV----ATTFLPTAS-NEATLRPSFVRDEDERPRVAYNQ----------FS	38
ZmF3H           -------MAPVSIS-AVPFLPTAAEGETNVRASFVREEDERPKVPHDR----------FS	42
AcFLS           --------MEVE---RVQAIATLTANLGTIPSEFIRSDHERPDLTTYH---------GPV	40
MusaFLS1        --------MEVE---RVQAIASLSVATNDIPPEFVRSEHEQPGITTYR---------GLV	40
MusaFLS3        --------MEVE---RVQVIASVCAADGVMPPEFIRSEHEQPGITTYR---------GPA	40
MusaFLS4        --------MEVE---RVQVIASVCAADGVMPPEFIRSEHEQPGITTYR---------GPA	40
MusaFLS2        --------MAQV---KVQAIASMLHSKDTIPPEFVRVEDEQPGATTYR---------GPA	40
AtFLS1          --------MEVE---RVQDISSSSLLTEAIPLEFIRSEKEQPAITTFR---------GPT	40
ZmFLS           --------MGGETHLSVQELAAS---LGALPPEFVRSEQDQPGATTYRG--------AAV	41
OsFLS           -------MAEVQ---SVQALASS---LAALPPEFVRSEHERPGATTFRG--------GDA	39
AtANS           -------MVAVE---RVESLAKS--GIISIPKEYIRPKEELESINDVFLEEKK----EDG	44
MusaANS         MATK--VVSVAP---RVEILAKS--GINEIPTEYVRPESERLDLGDAFEEVKKA---AEG	50
ZmANS           MESSPLLQLPAA---RVEALSLS—GLSAIPPEYVRPADERAGLGDAFDLARTHANDHTA	55
OsANS           -------MTDAE--LRVEALSLS—GASAIPPEYVRPEEERADLGDALELARAASDDDAT	49
                                .            :  .::*   :

AtF3H           DEIPVISLAGI--DDVD----GKRGEICRQIVEACENWGIFQVVDHGVDTNLVADMTRLA	89
MusaF3H1        LDIPVISLSGI--DDD--AAGGKRAEIRRKVVEACEDWGIFQVVDHGVDAGLISDMTRLA	91
MusaF3H2        SDIPVISLAGV--DDEDGATGGRRAEIRRKIVEACEDWGLFQVLDHGVDAGMISEMTRLA	93
OsF3H           DAVPVISLQGI--DE------AARAEIRARVAGACEEWGIFQVVDHGVDAGLVADMARLA	90
ZmF3H           DEVPVVSLEGI--DGP-----RRRAEIRGRVAAACEDWGIFQVVDHGVDAALVADMARLA	95
AcFLS           PELPVIDLANS-----------SQENVVKQISEAAREYGIFQLVNHGIPNEVINELQRVG	89
MusaFLS1        PEIPVIDLEDG-----------DEGRVTRAIAEASQEWGIFQLVNHGIPGEVIRALQRVG	89
MusaFLS3        PEIPVIDLAGA-----------DRDRLTIAVAEASREWGIFQLLNHGIPREVIRELQRVG	89
MusaFLS4        PEIPVIDLAGA-----------DRDRLTIAVAEASREWGIFQLLNHGIPREVIRELQRVG	89
MusaFLS2        PAVPVIDLADT-------------DRVVDAIADASREWGIFQLVNHGIPAAVIGELQRVG	87
AtFLS1          PAIPVVDLSDP-----------DEESVRRAVVKASEEWGLFQVVNHGIPTELIRRLQDVG	89
ZmFLS           PDAPVIDISEP--------------GFGARMAAAAREWGLFQVVNHGVPSAAVAELQRVG	87
OsFLS           PEIPVIDMAAP--------------ESAARVAEAAAEWGLFQVVNHGVPAAAVAELQRVG	85
AtANS           PQVPTIDLKNI--ESDD---EKIRENCIEELKKASLDWGVMHLINHGIPADLMERVKKAG	99
MusaANS         PQIPVVDLQGF--DSPD---EEVRRACVEEVRKAATEWGVMHIVNHGIPLELVEQLRKVG	105
ZmANS           PRIPVVDISPFLDSSSQ---QQQRDECVEAVRAAAADWGVMHIAGHGIPAELMDRLRAAG	112
OsANS           ARIPVVDISAF---DND---GDGRHACVEAVRAAAEEWGVMHIAGHGLPGDVLGRLRAAG	103
                   *.:.:                      :  *. ::*:::: .**:    :  :   .

AtF3H           RDFFALPPEDKLRF--DMSGGKKGGFIVSSHLQGEAVQDWREIVTYFSYPVRNRDYSRWP	147
MusaF3H1        KEFFALSPDEKLLF--DMSGGKKGGFIVSSHLQGEAVQDWREIVTYFSYPIRARDYSRWP	149
MusaF3H2        RDFFALPPEEKLRF--DMSGGKKGGFIVSSHLQGEAVQDWREIVTYFSYPIGARDYSRWP	151
OsF3H           RDFFALPPEDKLRF--DMSGGKKGGFIVSSHLQGEAVKDWREIVTYFSYPVKSRDYSRWP	148
ZmF3H           RDFFALPPEDKLRF--DMSGGKKGGFIVSSHLQGEAVQDWREIVTYFSYPVKARDYSRWP	153
AcFLS           KEFFQLPQEEKEVYATVPDSGSFEGYGTKLQKDLEGKKAWVDYLFHNVWPKHKINYKFWP	149
MusaFLS1        REFFELPPEEKEKYAAAP--GSLQGYGTKLQKDLEGKKAWVDFLFHNIWPPTHVDHRAWP	147
MusaFLS3        KQFFELPQEEKEMYAMECKPGSSEGYGTKLQRELEGKKAWVDFFFHYISPPARVNHAIWP	149
MusaFLS4        KEFFELPQEEKEMYAMEFKPGSSEGYGTKLQRELEGKKAWVDFFFHYVSPPARVNHAIWP	149
MusaFLS2        REFFELPQEEKESYAADPRSGSIEGYGTQIQKDPNGKKAWGDYLFHNVWPPSRINHGMWP	147
AtFLS1          RKFFELPSSEKESVAKPEDSKDIEGYGTKLQKDPEGKKAWVDHLFHRIWPPSCVNYRFWP	149
ZmFLS           RAFFALPTEEKERYAMDPASGKIEGYGTKLQRDLEGKKTWNDFFFHVVAPPEKVDHAVWP	147
OsFLS           REFFALPQEEKARYAMDASSGKMEGYGSKLQKDLEGKKAWADFFFHNVAPPAMVNHDIWP	145
AtANS           EEFFSLSVEEKEKYANDQATGKIQGYGSKLANNASGQLEWEDYFFHLAYPEEKRDLSIWP	159
MusaANS         KEFFDLPIEQKEQYANDQSSGKIQGYGSKLANNASGQLEWEDYFFHLIFPEEKINISIWP	165
ZmANS           TAFFALPVQDKEAYANDPAAGRLQGYGSRLATNTCGQREWEDYLFHLVHPDGLADHALWP	172
OsANS           EAFFALPIAEKEAYANDPAAGRLQGYGSKLAANASGKREWEDYLFHLVHPDHLADHSLWP	163
                  ** *.  :*             *:      :  .   * : . :   *    :   **

AtF3H           DKPEGWVKVTEEYSERLMSLACKLLEVLSEAM-GLEK-ESLTNAC----------VDMDQ	195
MusaF3H1        DKPEGWRSVVESYSEKLMGLACKLLEVLSEAM-GLDK-EALTEAC----------IDMDQ	197
MusaF3H2        DKPEGWRSVVESYSEKLMGLACKLLEVLSEAM-GLDK-EALTDAC----------VDMDQ	199
OsF3H           DKPAGWRAVVEQYSERLMGLACKLLGVLSEAM-GLDT-NALADAC----------VDMDQ	196
ZmF3H           DKPAAWRAVVERYSEQLMALACRLLGVLSEAM-GLDT-EALARAC----------VDMDQ	201
AcFLS           RNPPAYRKANEEYTKHLQVVVDKMHSYLSLGL-GLES-HVLKEAVGG--------DDLEY	199
MusaFLS1        ENPVDYRKANEEYAKHLVGLVEKMLVSLSKGL-GLEA-DVLKHAVGG--------DDLEF	197
MusaFLS3        KNPADYRKANEEYGKHLVGLVDKMLMTLSRGL-GLEE-HVLKGALGG--------DGLEL	199
MusaFLS4        KNPSDYRKANEEYAKHLVGLVDKMLTTLSRGL-GLEE-HVLKGALGG--------DGLEL	199
MusaFLS2        RQPSSYREANEEYTKYLLGILDKILDSLSLGL-GLEG-SALKKALGG--------EEMDM	197
AtFLS1          KNPPEYREVNEEYAVHVKKLSETLLGILSDGL-GLKR-DALKEGLGG--------EMAEY	199
ZmFLS           RSLAGYREANEEYCRHMQRLTRELFEHLSLGL-GLHG-GAMAEAFGG--------DGLVF	197
OsFLS           SHPAGYREANEEYCKHMQRLARKLFEHLSTAL-GLDG-GAMWEAFGG--------DELVF	195
AtANS           KTPSDYIEATSEYAKCLRLLATKVFKALSVGL-GLEP-DRLEKEVGGL-------EELLL	210
MusaANS         KQPTDYIEVTKKFGRQLRAVVTKMLEVLSLGL-GLEE-GKLDRELGGM-------EDLLM	216
ZmANS           AYPPDYIAATRDFGRRTRDLASTLLAILSMGLLGTDRGDALEKALTTTTTRTAADDDLLL	232
OsANS           ANPPEYVPVSRDFGGRVRTLASKLLAILSLGL-GLPE-ETLERRLRGHEL-AGVDDDLLL	220
                     :  .   :      :   :   ** .: *      :

AtF3H           KIVVNYYPKCPQPDLTLGLKRHTDPGTITLLLQDQVGGLQATRDNGKTWITVQPVEGAFV	255
MusaF3H1        KVVVNYYPKCPQPDLTLGLKRHTDPGTITLLLQDQVGGLQATKDGGKTWITVQPVEGAFV	257
MusaF3H2        KVVVNYYPMCPQPDLTLGLKRHTDPGTITLLLQDQVGGLQATKDGGKTWITVQPVEGAFV	259
OsF3H           KVVVNFYPKCPQPDLTLGLKRHTDPGTITLLLQDLVGGLQATRDAGKTWITVQPIPGSFV	256
ZmF3H           KVVVNFYPRCPQPDLTLGLKRHTDPGTITLLLQDLVGGLQATRDGGRTWITVQPVEGAFV	261
AcFLS           LLKINYYPPCPRPNLALGVVAHTDMSSLTILVPNEVPGLQVFKDD--HWFDAKYIPNALI	257
MusaFLS1        LLKINYYPPCPRPDLALGVVAHTDMSAITILIPNDVPGLQVFKDD--HWFDAKYVPDAII	255
MusaFLS3        RLKMNYYPPCPRPDLALGVVAHTDMCAITFLVPNLVPGLQVFKDE--HWIDVNFIPNAVI	257
MusaFLS4        RLKMNYYPPCPRPDLALGVVAHTDMCAITFLVPNLVPGLQVFKDE--HWIDVNFIPNAVI	257
MusaFLS2        LLKINYYPPCPRPDLALGVVAHSDLSAVTILVPSDVPGLQISKDD--RWIDIDYVPGALI	255
AtFLS1          MMKINYYPPCPRPDLALGVPAHTDLSGITLLVPNEVPGLQVFKDD--HWFDAEYIPSAVI	257
ZmFLS           LQKINFYPPCPQPELTLGVAPHTDMSTLTVLVPNEVQGLQVFKDG--QWYEAKYVPDALI	255
OsFLS           LHKINFYPPCPEPELTLGVAPHTDMSTFTVLVPNDVQGLQVFKDG--HWYDVKYVPDALI	253
AtANS           QMKINYYPKCPQPELALGVEAHTDVSALTFILHNMVPGLQLFYEG--KWVTAKCVPDSIV	286
MusaANS         QLKINYYPICPQPDLALGVEAHTDISALSFILHNMVPGLQIYYGG--RWVTAKCVPDSII	274
ZmANS           QLKINYYPRCPQPELAVGVEAHTDVSALSFILHNGVPGLQVLHGA--RWVTARHEPGTII	290
OsANS           QLKINYYPRCPRPDLAVGVEAHTDVSALSFILHNGVPGLQVHHAG--SWVTARPEPGTIV	278
                   :*:** **.*:*::*:  *:*   .:.:: . * ***        *       .:.:

AtF3H           VNLGDHGH-----------FLSNGRFKNADHQAVVNSNSSRLSIATFQNPAPDAT-VYPL	303
MusaF3H1        VNLGDHGH-----------FLSNGRFKNADHQAVVNSNCSRLSIATFQNPAPEAI-VYPL	305
MusaF3H2        VNLGDHGH-----------YLSNGRFKNADHQAVVNSSCSRLSIATFQNPAPEAI-VYPL	307
OsF3H           VNLGDHAHIMHLLGNVNLQYLSNGRFKNADHQAVVNSDCCRLSIATFQNPAPDAM-VYPL	315
ZmF3H           VNLGDHGH-----------LLSNGRFKNADHQAVVNSECSRLSIATFQNPAPDAT-VYPL	309
AcFLS           CHIGDQLE-----------ILSNGKYKSVLHRTTVNKEKSRMSWPVFCSPPGDTM-IGPL	305
MusaFLS1        VHIGDQIE-----------KLSNGRYKSVLHRTTVNKEKARMSWPVFCSPPGETV-IGPL	303
MusaFLS3        VHIGDQIE-----------ILSNGTYKSVLHRTTVNKEKVRMSWPVFCAPPGEMV-IGPL	305
MusaFLS4        VHIGDQIE-----------ILSNGTYKSVLHRTTVNKEKVRMSWPVFCAPPGEMV-IGPL	305
MusaFLS2        IHIGDQIE-----------ILSNGKYKSVLHRATVNKEKARISWPVFCSPPPEMT-VGPL	303
AtFLS1          VHIGDQIL-----------RLSNGRYKNVLHRTTVDKEKTRMSWPVFLEPPREKI-VGPL	305
ZmFLS           VHIGDQIE-----------IFSNGAYKAVLHRTTVNKEKTRMSWPMFVEPPGELV-VGPH	303
OsFLS           IHIGDQIE-----------ILSNGRYKAVLHRTTVDKDRTRMSWPVFVEPPPEHV-VGPH	301
AtANS           MHIGDTLE-----------ILSNGKYKSILHRGLVNKEKVRISWAVFCEPPKDKIVLKPL	317
MusaANS         MHVGDCLE-----------ILSNGQYKSILHRGLVNKEKVRISWAVFCEPPKDKIVLKPL	323
ZmANS           VHVGDALE-----------ILSNGRYTSVLHRGLVNREAVRISWVVFCEPPPDSVLLHPL	339
OsANS           VHVGDALE-----------ILTNGRYTSVLHRGLVSRDAVRLSWVVFCEPPPESVLLQPV	327
                 ::**               ::** :.   *:  *. .  *:*   *  *. :   : *

AtF3H           K--VREGEKAILEEPITFAEMYKRKMGRDLELARLKKLAKEERDHK-EVDKP--------	352
MusaF3H1        A--VREGEKPILDEPITFAEMYRRKMSRDLELAKLKKLAKTEQQQKPELLEK-------T	356
MusaF3H2        A--IREGEKPILDEPITFSEMYRRKMSRDLKLANLKKLAKAEHQRL-ELPEN-------A	357
OsF3H           A--VRDGEEPILEEPITFAEMYRRKMARDLELAKLKKKAKEQRQLQ-QAALPPPPPTQVA	372
ZmF3H           A--VRDGEAPILDHPITFAEMYRRKMARDIELARLKKQAKADKKQQ-QQSAN--------	358
AcFLS           PQLVNDEN-PPKFKTKKYKDYAYCKINK---------LPQ--------------------	335
MusaFLS1        PQLVSDEQ-PAQYKTKKYKDYAFCKLNK---------LPQ--------------------	333
MusaFLS3        QQLVGDES-PAKYKPKKYKDYAYCKLNK---------LPQ--------------------	335
MusaFLS4        QQLVGDES-PAKYKPKKYKDYAYCKLNK---------LPQ--------------------	335
MusaFLS2        PQFVSDQN-PAKYKTKKYKDYQYCKLNK---------LPQ--------------------	333
AtFLS1          PELTGDDN-PPKFKPFAFKDYSYRKLNK---------LPLD-------------------	336
ZmFLS           PKLVTEES-PAKYKAKKYKDYQHCKINK---------LPM--------------------	333
OsFLS           PQLVTDGS-PAKYKAKKFKDYRHCKINK---------LPM--------------------	331
AtANS           PEMVSVES-PAKFPPRTFAQHIEHKLFG---------KEQEELVSE-KN-----------	355
MusaANS         EELVADGT-PAKFPPRTFEQHIQHKLFK---------KTRGDFKTP--------------	359
ZmANS           PELVTEGH-PARFTPRTFKQHLDRKLFK---------KKQQHKAKA-EEEDGG------N	382
OsANS           QELLADGAGKPLFAPRTFKQHVQRKLFKKL-------KDQQDNNAA-AASNG--------	371
                              .  : :    *:

AtF3H           ----------VDQIFA	358
MusaF3H1        KDINLAKATGLDEILA	372
MusaF3H2        NDINVAKAKCLNEILA	373
OsF3H           AELAAQKPKSLDEILA	388
ZmF3H           KEFADSKP--LDAIFA	372
AcFLS           ----------------	335
MusaFLS1        ----------------	333
MusaFLS3        ----------------	335
MusaFLS4        ----------------	335
MusaFLS2        ----------------	333
AtFLS1          ----------------	336
ZmFLS           ----------------	333
OsFLS           ----------------	331
AtANS           ---------------D	356
MusaANS         ---------------N	360
ZmANS           GDHHRHEP---PPQTN	395
OsANS           ------------MITK	375
```

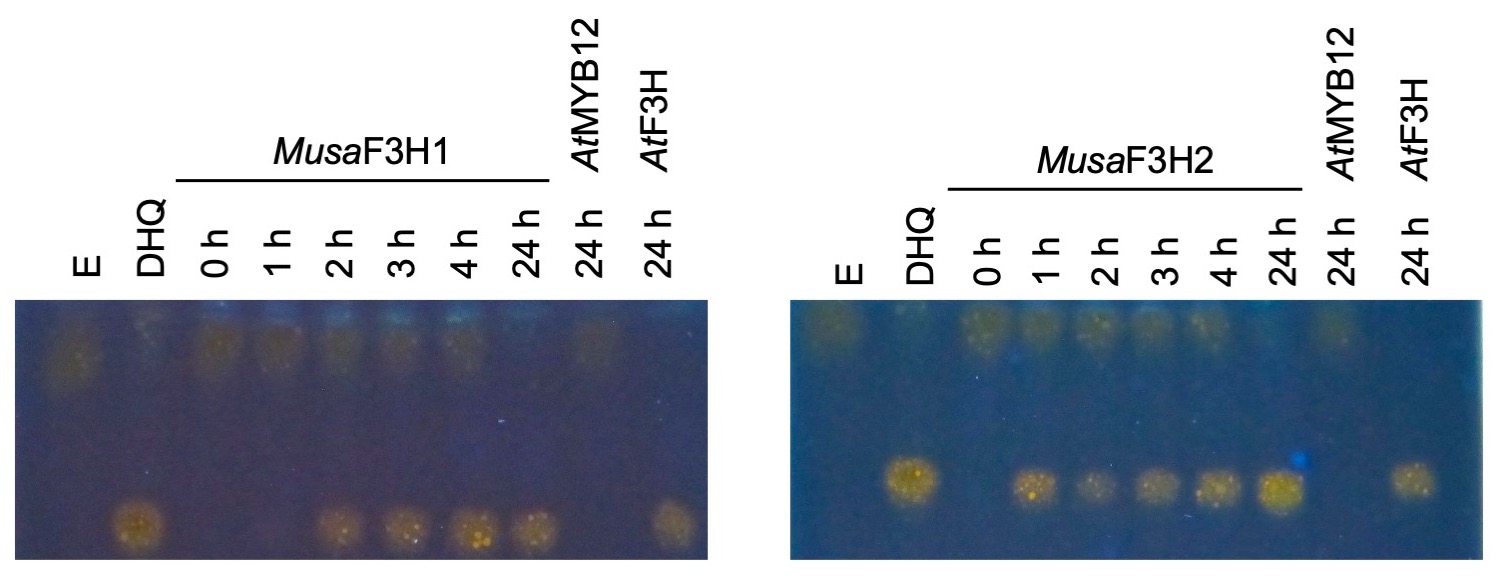

**File S3: *Musa*F3H1 and *Musa*F3H2 convert eriodictyol to dihydroquercetin.**

HPTLC analysis of F3H bioconversion assays using extracts from *E. coli* expressing recombinant *Musa*F3H1 or *Musa*F3H2. The substrate eriodictyol (E) and the product dihydroquercetin (DHQ) were used as standards. *At*F3H was used as a positive control. *At*MYB12 was used as a negative control.

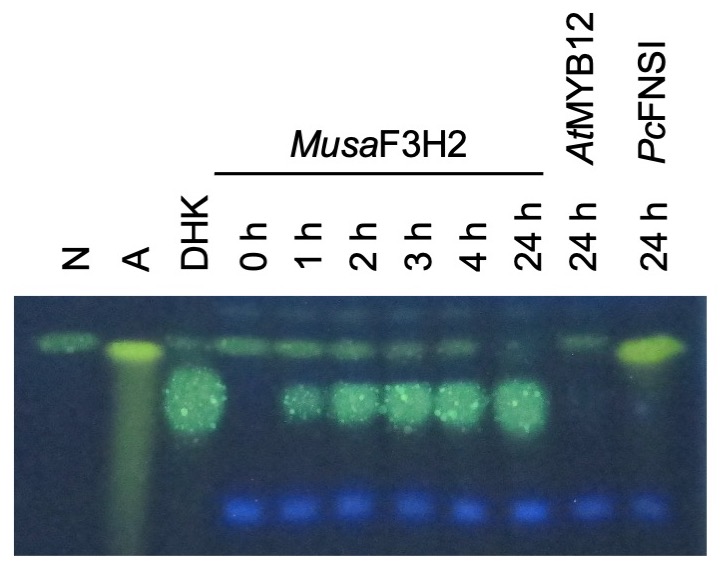

**File S4: Analysis of FNSI side activity of *Musa*F3H2.**

HPTLC analysis of FNSI bioconversion assay using extracts from *E. coli* expressing recombinant *Musa*F3H2. The substrate naringenin (N) and the product apigenin (A), as well as the F3H product dihydrokaempferol (DHK) were used as standards. *Pc*FNSI was used as a positive control. *At*MYB12 was used as a negative control.

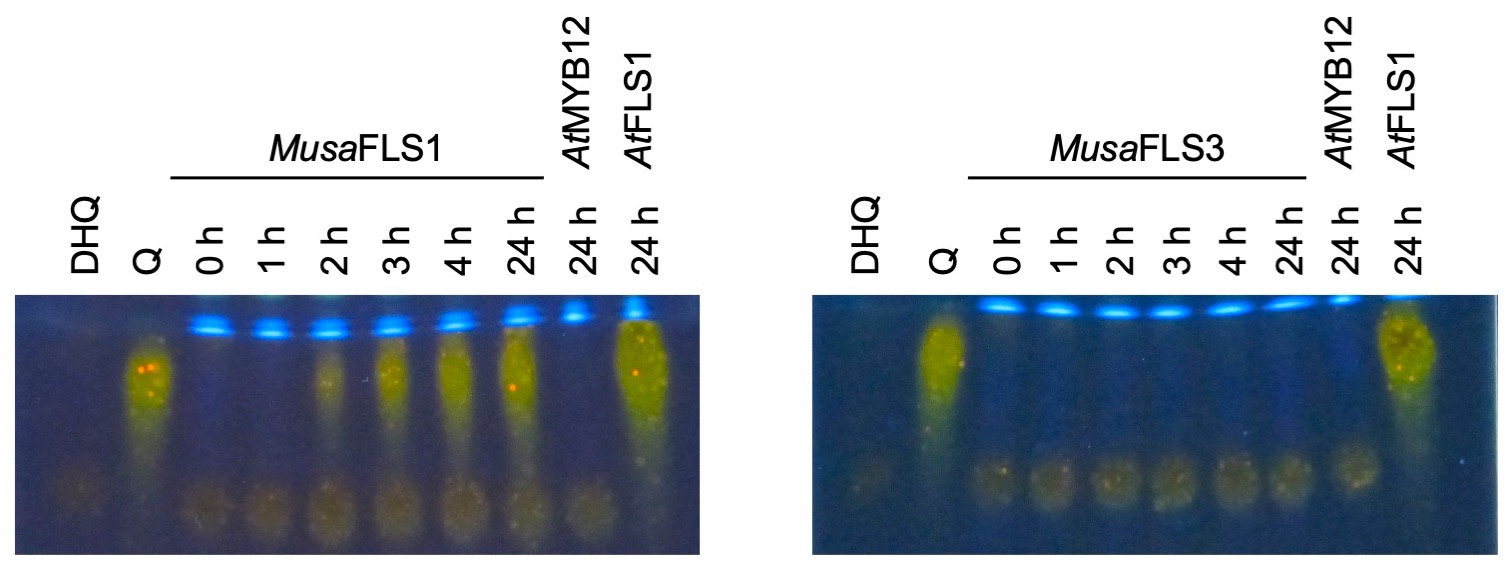

**File S5: MusaFLS1 converts dihydroquercetin to quercetin.**

HPTLC analysis of a FLS bioconversion assays using extracts from *E. coli* expressing recombinant *Musa*FLS1 or *Musa*FLS3. The substrate dihydroquercetin (DHQ) and the product quercetin (Q) were used as standards. *At*FLS1 served as a positive control and *At*MYB12 was used as a negative control.

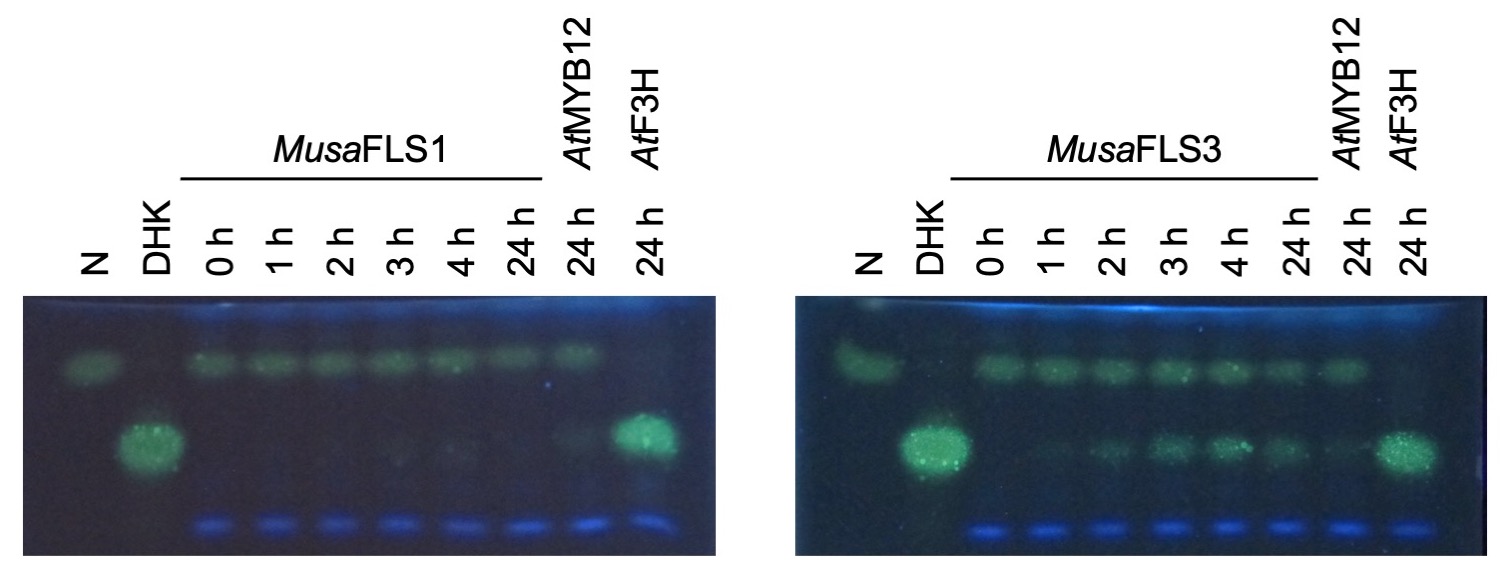

**File S6: Analysis of F3H side activity of *Musa*FLS1 and *Musa*FLS3.**

HPTLC analysis of F3H bioconversion assay using extracts from *E. coli* expressing recombinant *Musa*FLS1 or *Musa*FLS3. The substrate naringenin (N) and the product dihydrokaempferol (DHK) were used as standards. *At*F3H was used as a positive control. *At*MYB12 was used as a negative control.

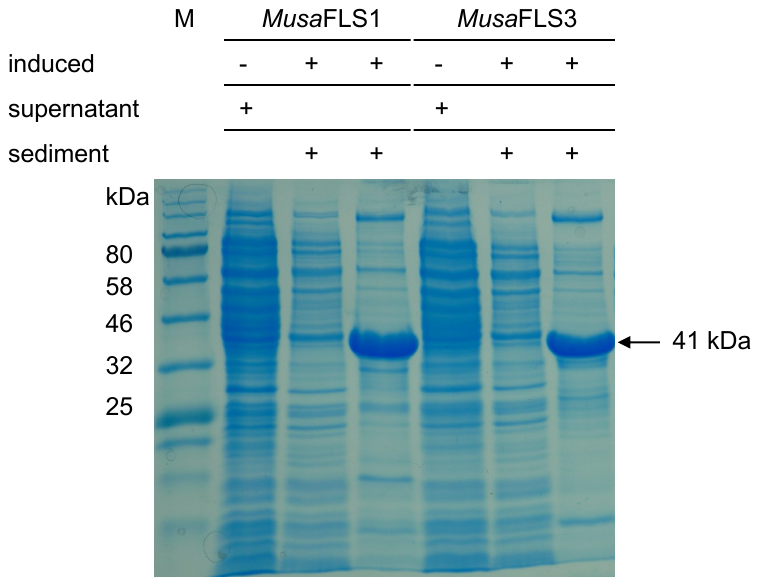

**File S7: Analysis of recombinant *Musa*FLS1 and *Musa*FLS3 in *E. coli* by SDS-PAGE.**

Sample separation was performed by SDS-PAGE (Laemmli, 1970) using a 4 % stacking gel and 12 % resolving gel. Protein staining was performed using colloidal Coomassie Brilliant Blue G-250 (Dyballa and Metzger, 2012). The first lanes for each sample shows the supernatant of *E. coli* cultures without addition of inducing arabinose, the second ones show the supernatant of induced *E. coli* cultures and the third lanes show the sediment of induced *E. coli* cultures. Including the polyhistidine-tag, *Musa*FLS1 and *Musa*FLS3 have a mass of about 41 kDa. The *Color Prestained Protein Standard, Broad Range* (NEB) was used as a standard.
